# High frequency bursts facilitate fast communication for human spatial attention

**DOI:** 10.1101/2024.09.11.612548

**Authors:** Kianoush Banaie Boroujeni, Randolph F. Helfrich, Ian C. Fiebelkorn, J. Nicole Bentley, Peter Brunner, Jack J. Lin, Robert T. Knight, Sabine Kastner

**Affiliations:** Princeton Neuroscience Institute, Princeton University, Princeton, NJ 08544; Departments of Psychology and Neurology and the Wu Tsai Institute, Yale University, New Haven, CT 06510, USA; Department of Neuroscience and Del Monte Institute for Neuroscience, University of Rochester, Rochester, NY 14627, USA; Department of Neurosurgery, University of Alabama at Birmingham, Birmingham, Alabama, USA; Department of Neurosurgery, Washington University in St Louis, St Louis, MO, USA; Department of Neurology and Center for Mind and Brain, University of California Davis, Davis, CA, USA; Departments of Psychology and Neuroscience, University of California Berkeley, Berkeley, CA, USA; Department of Psychology, Princeton University, Princeton, NJ 08544

**Author notes:** Corresponding author: Kianoush Banaie Boroujeni.

## Abstract

Brain-wide communication supporting flexible behavior requires coordination between sensory and associative regions but how brain networks route sensory information at fast timescales to guide action remains unclear. Using human intracranial electrophysiology and spiking neural networks during spatial attention tasks, where participants detected targets at cued locations, we show that high-frequency activity bursts (HFAb) mark temporal windows of elevated population firing that enable fast, long-range communications. HFAbs were evoked by sensory cues and targets, dynamically coupled to low-frequency rhythms. Notably, both the strength of cue-evoked HFAbs and their decoupling from slow rhythms predicted behavioral accuracy. HFAbs synchronized across the brain, revealing distinct cue- and target-activated subnetworks. These subnetworks exhibited lead-lag dynamics following target onset, with cue-activated subnetworks preceding target-activated subnetworks when cues were informative. Computational modeling suggested that HFAbs reflect transitions to population spiking, denoting temporal windows for network communications supporting attentional performance. These findings establish HFAbs as signatures of population state transitions, supporting information routing across distributed brain networks.

## Main text

Prioritizing information from the external environment to guide ongoing behavior and upcoming actions requires fast coordination of neural activity in large-scale networks distributed across distant brain areas.^1–6^ This coordination allows information to be routed selectively from sensory to higher level executive brain networks.^7–9^ Previous research, particularly in non-human primates, has shown that selective information routing emerges through dynamically changing neuronal interactions.^2,10,11^ Such studies highlight the role of oscillatory dynamics and transient changes in inter-areal coherence in enabling attentional selection and the flexible reconfiguration of neural pathways according to task demands. Yet, most prior investigations have focused on pairwise interactions between a few brain areas, leaving open the question of how fast neural dynamics emerge and enable flexible, large-scale information routing across distributed networks.

Investigating these questions in the human brain is challenging due to spatial or temporal constraints of available techniques. Studies examining network-level interactions have largely been based on connectivity maps derived from functional magnetic resonance imaging (fMRI). While fMRI studies provide valuable insights into functional networks, the temporal resolution cannot capture sub-second routing dynamics in attention tasks.^12–15^ Electroencephalography (EEG) and magnetoencephalography (MEG) offer high temporal resolution but have limited spatial resolution.^16,17^ Lastly, single unit recordings provide both fine temporal and spatial signals but lack the broad coverage for addressing questions of brain-wide network communication.^2,6,18^ Human intracranial electroencephalography (iEEG), offers a unique opportunity to address these challenges by providing spatially localized and temporally precise neural signals from multiple brain regions.^19,20^ High-frequency activity detected in iEEG signals correlates with different cognitive functions, including attention,^19–24^ and has been reported to index aggregated spiking activity, dendritic post-synaptic activity, state transitions into spiking regimes, or spike current leakage to local field potentials (LFPs).^25–29^ Additionally, high-frequency activities show long-range phase synchronization with different frequency bands, making them candidates for studying fast brain-wide communications.^21,30–33^ However, iEEG studies of attention have predominantly focused on fast-slow interactions, such as cross-frequency coupling between high- and low-frequency rhythms, which may not fully capture the dynamics of fast-fast interactions between high-frequency activities.^21,30,34–37^

Here, we address this gap by identifying transient high-frequency activity bursts (HFAbs) in human iEEG data from epilepsy patients performing spatial attention tasks, hypothesizing that HFAbs reflect periods when neural populations establish temporal alignment across distributed regions, creating windows for selective information routing. HFAbs during sensory cue processing predicted successful detection of upcoming targets. They were locally phase-locked to slow rhythms (4-25 Hz), and transiently decoupled during cue and target processing, with decoupling associated with correct performance. Across brain networks, HFAbs temporal relationships were largely characterized by zero-lag synchronization, identifying functionally specialized subnetworks with distinct topographical distributions and temporal activation sequences. Specifically, HFAbs in cue-subnetwork preceded target-subnetwork activity following target onset, when sensory cues provided relevant information about the target location. Using computational modeling, we found that HFAbs mark state transitions in neural populations into a spiking state, characterized by bouts of elevated neuronal firing, indicating enhanced population excitability enabling synchronized and temporally ordered activation patterns across regions.

## Results

### HFAbs track spatial attention and predict behavioral accuracy

We analyzed iEEG data from two spatial attention tasks in which patients were cued either exogenously or endogenously to a spatial location to detect visual targets (**Fig. 1a**, **Extended Data Fig. 1a**). Throughout the paper, the main figures focus on the results of experiment 1, while the results of experiment 2 are presented as supplementary information, unless mentioned otherwise (for brain group-average heatmap plots, we combined both experiments to improve 3D rendering coverage).

**Fig. 1.**
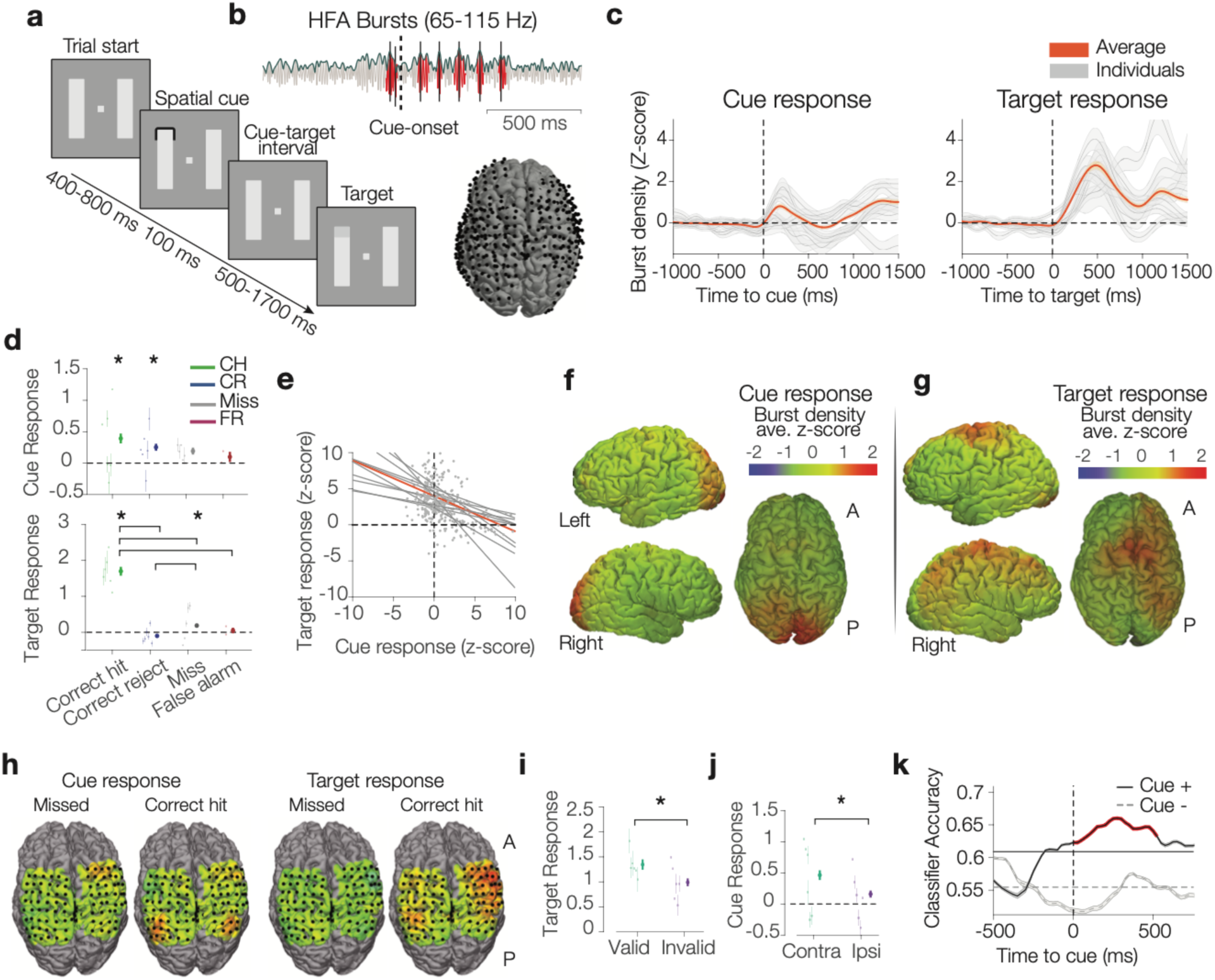
HFAb activation profile predicts behavioral outcome on a trial-by-trial basis. **a,** Experiment 1 task outline: two bars appear on the screen, followed by a transient cue indicating the likely target location. After a variable delay, a near-threshold contrast change appears at the cued or, less often, at an equidistant uncued location. In 10% of trials no target appears. Subjects report the change detection. **b**, An example trial showing detected high-frequency bursts (HFAbs, in red) around the cue onset. The brain shows localization of electrodes across all subjects in experiment 1. **c**, Normalized HFAb density for individual subjects and the group average, aligned to the cue (left) and target onset (right); shaded error bars denote mean ± SEM. **d**, HFAb responses to cues (top) and targets (bottom) grouped by trial outcome (n=6, median ± SEM, each line represents one subject, with thick lines showing group average, horizontal lines showing significant differences between groups (Dunn’s test, *p* < 0.05), and asterisks denoting non-zero responses, Wilcoxon signed-rank test, *: *p* < 0.001). **e**, Correlation of HFAb responses to cue and target, for channels with significant activation to either or both cue and target (each point represents one channel, each line represents regression for one subject, the orange line shows regression across all channels). (**e, g**) Group average heatmap 3D rendering of HFAb responses to (**a**) cues and (**g**) targets for correct trials. **h**, An individual subject example of HFAb responses to cue and target, for correct and incorrect trials (black circles indicate electrode locations). **(i,j)**, HFAb responses to (**i,** GLME, *t* = 4.7, *: *p* < 0.001) targets for valid and invalid cue conditions and (**j**, GLME, *t* = 5.1, *: *p* < 0.001) to cues ipsi- and contra-lateral to electrodes (n=6, median ± SEM). **k**, Classifier accuracy in predicting trial outcome based on HFAb density around the cue onset for cue responsive electrodes and cue unresponsive electrodes (error bars: SEM across realizations and cross validations). Red lines indicate points significantly above baseline and chance (two-sided binomial test, *p* < 0.05, FDR-corrected for multiple dependent comparisons).

In experiment 1, patients performed a spatial attention task.^23^ Each trial started with the presentation of two vertical or horizontal bar stimuli. Patients were instructed to fixate their gaze at the center of the display. A transient spatial cue appeared at the end of one bar, exogenously cueing an upcoming target location. Following a delay period, a target (i.e., luminance changes at perceptual threshold) appeared at the cued location or infrequently at equally distant non-cued locations. Patients responded when they detected the target (**Fig. 1a**). In experiment 2, patients were endogenously cued to a hemifield and reported a target if it appeared in the cued hemifield^3^ (**Extended Data Fig. 1a**, see **Methods,** see **Extended Data Fig. 1b** for detailed performance breakdowns).

First, we used an adaptive method^6^ and detected bursts of high frequency activity at each electrode (HFAbs, 65-115 Hz, intermittent high amplitude events lasting more than 2.5 frequency cycles, average burst length 36.2 ms; **Fig. 1b**). The frequency band was selected based on average spectral peaks observed across subjects (91.2 ± 20.9 Hz, n = 12, see **Methods**). HFAbs occurred frequently (0.1 ± 0.01 Hz) on average in our data, approximately 15-fold higher than recently described cortical ripples (0.006 ± 0.0008 Hz),^33^ suggesting distinct characteristics of HFAbs despite overlapping spectral properties (see **Methods**). We then calculated the HFAb density (HFAb events per unit of time) for each electrode to examine their evoked response during different task epochs. HFAbs showed higher density averaged across all electrodes in response to both cue and target (**Fig. 1c**). We measured HFAb responses following cue and target onsets across different trial outcomes (hit, reject, miss, and false alarm). At the population level, HFAbs activated to cue onsets in correct hit and reject trials (Wilcoxon signed-rank test, *p* < 0.001) and differed significantly from missed and false alarm trials (Kruskal-Wallis test, *p* = 0.001, Dunn’s test, *p* = 0.008 and *p* = 0.01, respectively; **Fig. 1d**). Target responses were also enhanced in correct hit trials compared to other outcome conditions (Kruskal-Wallis test, *p* < 0.001; Dunn’s test, *p* < 0.001; **Fig. 1d**). Similar results were observed in experiment 2 (**Extended Data Fig. 1c,d**). Overall, 10.7 ± 2% of channels (n = 12) showed significant activation to cues, whereas 33.2 ± 3% of channels (n = 12) were responsive to targets (Wilcoxon signed-rank test, *p* < 0.05). The broader activation to targets is in accord with greater complexity of target processing, engaging brain regions involved in sensory processing, motor planning, and decision-making. Among channels that responded to cues and/or targets, HFAb responses to targets were negatively correlated with those to the cue (Spearman correlation, *p* < 0.001, *r* = −0.36, **Fig. 1e**), suggesting distinct electrode populations process cues and targets. Cue responses were prominent in occipital and parietal regions, including areas in extrastriate cortex, intraparietal sulcus (IPS), temporoparietal junction (TPJ), superior parietal lobule (SPL), and inferior parietal lobule (IPL) (**Fig. 1f**), whereas target responses were more widely distributed including superior, middle, and inferior frontal gyrus, precentral and postcentral gyrus, IPL, SPL, and TPJ (**Fig. 1g**). This activation profile was evident only for correct trials both for cue- and target responses (**Extended Data Fig. 1e,f**). It is worth noting that for experiment 2, we also observed enhanced cue responses in the frontal gyrus consistent with more top-down attentional control in endogenous cueing. See **Extended Data Table 1** for detailed electrode positions of the cue.

We investigated the effect of cue validity (targets at cued vs uncued location) and laterality (visual field ipsilateral versus contralateral to electrodes) on cue and target responses using a Generalized Linear Mixed Effect (GLME) model (see **Methods**). We found a main effect of outcome on both cue response (*t* = 2.8, *p* = 0.005) and target response (*t* = 15.2, *p* < 0.001), a main effect of validity on the target response (**Fig. 1i**, *t* = 4.7, *p* < 0.001), and a main effect of laterality on the cue response (**Fig. 1j**, *t* = 5.1, *p* < 0.001). No significant main effect was observed for laterality on target (***p*** = 0.13), or validity on cue responses (**Extended Data Fig. 1g**, *p* = 0.32, See **Extended Data Fig. 1h** for experiment 2).

Given that HFAb activation to sensory cues was associated with outcome accuracy, we examined if HFAb density following cue onset in cue responsive electrodes (Cue+) predicted correct trials. A classifier was trained on the grand mean of HFAb density across selected electrodes using a 350 ms sliding window around the cue onset, with trial-level outputs labeled as correct or incorrect (see **Methods)**. In Cue+ electrodes, HFAb density within 500 ms post-cue predicted correct performance above baseline and chance levels (binomial test, FDR corrected for dependent samples, *p* < 0.05, **Fig. 1k**). This prediction was consistent across both experiments (**Extended Data Fig. 1i**) and was not dependent on an individual subject.

These findings indicate that HFAbs occur frequently in response to sensory cues and targets and exhibit distinct spatial profiles across the brain. HFAbs predicted performance accuracy following cues on a trial-by-trial basis. Together, these spatially selective and behaviorally predictive responses support HFAbs as neural signatures of attention-related processing.

### HFAbs are coupled to slow rhythms and decouple in response to cues and targets

High-frequency activity dynamics in brain networks are linked to low-frequency dynamics, especially through phase-locking relationships with theta rhythms (4-8 Hz).^30^ We asked whether HFAbs synchronized to the phase of low-frequency rhythms, and whether these cross-frequency dynamics were associated with task variables. For each electrode, we extracted the LFP around the HFAb centers and measured both the HFAb-triggered LFP average and the phase locking value (PLV) of HFAb peaks to low-frequency LFP dynamics. Spectral analysis revealed HFAb-triggered spectral peaks (**Fig. 2a**, top) and phase locking peaks (**Fig. 2a**, bottom) in theta (4–8 Hz), alpha (8–14 Hz), and beta (15–25 Hz) frequency bands. Phase-locking peaks occurred at one or more narrow frequency bands, showing different mean phases (**Fig. 2b**; see **Extended Data Fig. 2** for additional single-electrode examples, mean phase distributions in theta/alpha and beta bands across subjects, and spectral and phase-locking peak details). Phase-locking was evident in most electrodes across all subjects (reliable peaks in 97.8% ± 0.8% of electrodes; mean peak frequency 9 ± 0.24 Hz; significant peaks marked by black dots, **Fig. 2c**; see **Extended Data Fig. 3a,b** for topography of phase-locking peak frequencies).

**Fig. 2.**
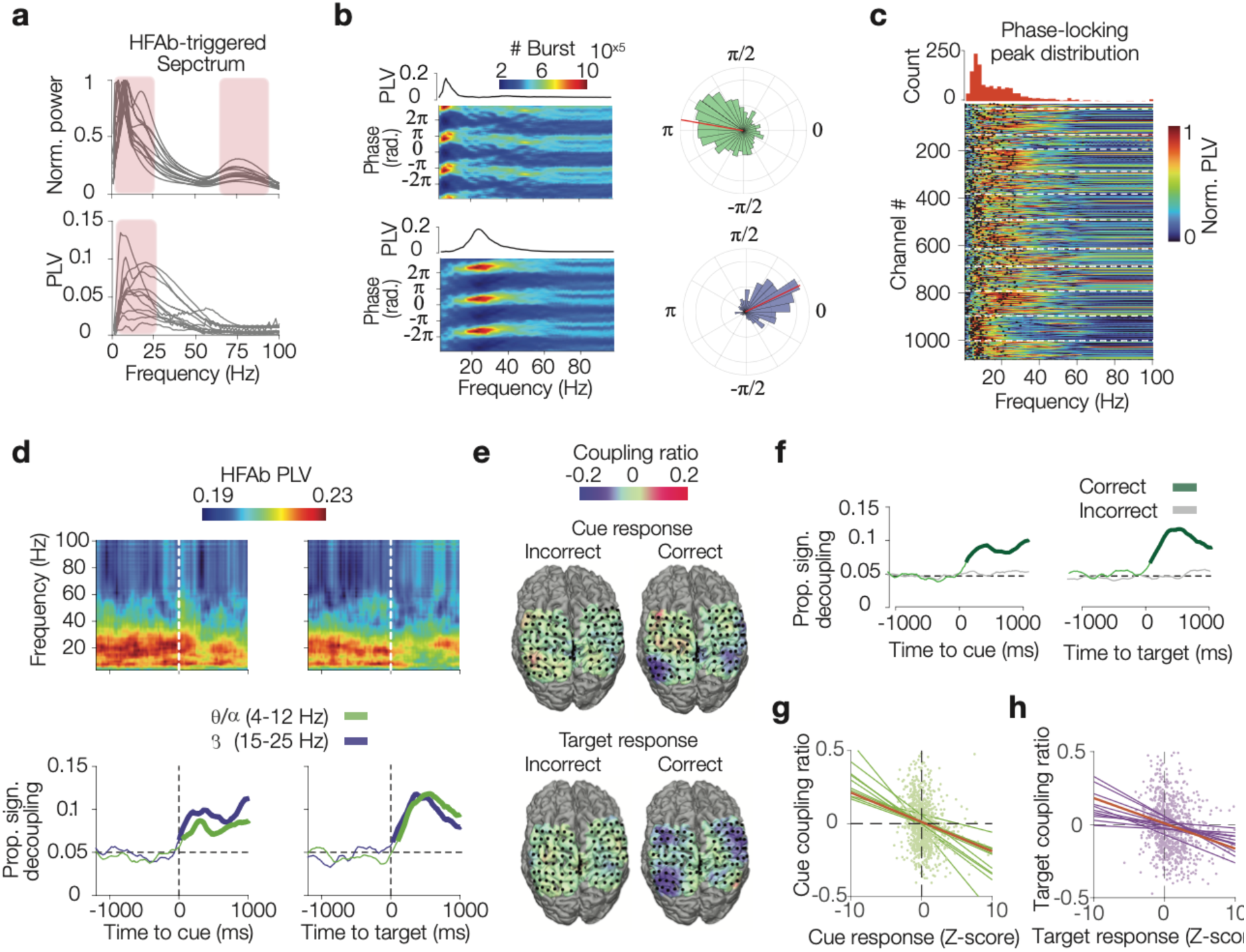
HFAbs dynamically phase lock to low-frequency LFPs and decouple transiently after cue and target onsets. **a**, HFAb-triggered LFP spectrum (top) and phase locking values (PLV, bottom) across all electrodes in both experiments (each line represents one subject). **b**, Examples of phase-frequency distribution of HFAbs for an electrode with theta phase locking (top) and beta phase locking (bottom). Polar histograms of phase distributions at the phase-locking frequencies are shown in the right-hand panels. **c**, PLVs for individual electrodes (each row). Black dots indicate the maximum peak of PLV for each electrode, and white dashed lines indicate subject boundaries. Phase-locking peak distribution is shown above. **d**, Top: time-resolved PLV analysis averaged across all subjects aligned to cue (left) and target (right) onsets in experiment 1. Bottom: The proportion of time points where phase locking dropped significantly below the baseline averaged across all electrodes (two-sided random permutation test, *p* < 0.05). Thick lines indicate segments significantly different from the chance level (two-sided binomial test, *p* < 0.05, FDR-corrected for multiple dependent comparisons; see **Methods**). **e**, Example subject brain heatmap showing coupling ratio between HFAb and low frequency (4-25 Hz) LFPs after cue (top) and target onsets (bottom) for correct and incorrect trials. **f**, Proportion of time points with significant decoupling from low-frequency LFPs (4-25 Hz) after cue and target onsets for correct and incorrect trials. **g, h**, Regression plots showing correlation of coupling ratios following cue onset with cue responses (green, **g**), and coupling ratios following target onset with target responses (purple, **h**). Scatter points denote electrodes, lines indicate individual subjects with orange line showing the regression across all subjects.

To test whether sensory cues or target processing affected HFAb phase-locking, we analyzed time-resolved PLV in sliding windows around cue and target onsets (HFAb count controlled per window; see **Methods**). HFAbs showed a transient decrease in their phase-locking (decoupling) in theta/alpha and beta frequency bands following cue and target onsets (**Fig. 2d, Extended Data Fig. 3c,** randomization test, *p* < 0.05, FDR corrected for dependent samples). This decoupling was not attributable to event-triggered potentials or the measurement approach (**Extended Data Fig. 3d,e** see **Methods**). Furthermore, changes in HFAb coupling strength were only observed in correct trials, not in incorrect trials (**Fig. 2e,f**, HFAb counts controlled across trial conditions, see **Methods**, see **Extended Data Fig. 3f** for individual examples). Similar results were observed in experiment 2 (**Extended Data Fig. 3g**).

Next, we examined whether changes in electrodes’ HFAb coupling to low frequencies were related to their responsiveness to cues and targets. For each subject, we correlated coupling ratios (changes in HFAb coupling to <25 Hz after cues or targets relative to baseline; see **Methods**) with burst density. Coupling ratios at cue and target onsets were negatively correlated with burst density following these events suggesting that electrodes with strong HFAb responses also showed decoupling (**Fig. 2g,h**, Spearman correlation, *r* = −0.22, *p* < 0.001, and *r* = −23, *p*<0.001, respectively). GLME models showed a main effect of cue response on coupling ratio after cue onset (**Fig. 2g** *t* = −6.17, *p* < 0.001), and of target response after target onset (**Fig. 2h**, *t* = −5.05, *p* < 0.001). No significant cross-effects were observed (cue response after target onset: *p* = 0.15; target response after cue onset: *p* = 0.21; **Extended Data Fig. 3h,i**).

Overall, HFAbs were predominantly phase-locked to theta, alpha, and beta rhythms. However, their coupling strength to slower rhythms decreased during cue and target processing, specifically in correct trials. This selective decoupling during correct trials suggests that releasing HFAbs from their default phase-locked state may facilitate the transition from internally regulated network activity to externally driven sensory processing required for successful task performance.

### HFAbs synchronized brain-wide and their network-level synchronization identified functionally specialized subnetworks

We found that HFAbs evoked by cue and target demonstrated different topographical distributions, largely non-overlapping responsive electrode populations. Additionally, HFAbs transiently decoupled from local low-frequency rhythms during task events, suggesting reduced local rhythmic constraints during active processing. The distinct spatial patterns and release from local phase-locking raise the possibility that HFAbs may instead synchronize with HFA in distant regions processing similar information. We therefore asked whether brain networks could be divided to distinct subnetworks based on the temporal relationships of their HFAbs, and whether subnetworks with similar synchronization patterns are functionally specialized.

We considered HFAbs outside the cue/delay and target/response periods to avoid stimulus-driven synchronization effects and measured the power of high frequency activity (65-175 Hz, HFA, to capture broader spectral contents) in one electrode aligned to the center of HFAb events in another electrode (HFAb-triggered HFA). HFAb-triggered HFA between electrodes showed synchronized high-frequency amplitude increases (**Fig. 3a**, similar results were observed for HFAb-HFAb). These synchronized patterns exhibited rhythmic modulation at low-frequency rhythms, with spectral peaks predominantly in the theta band (4-8 Hz) (**Fig. 3b**, top). These patterns were consistent across subjects and experiments (**Fig. 3b**, bottom, **Extended Data Fig. 4a,b**). HFAb synchronization strength (the sharpness of HFAb-triggered HFA) was inversely related to inter-electrode distance, with closer sites showing stronger synchronization (**Fig. 3c**, **Extended Data Fig. 4c**, Spearman correlation between HFAb-triggered HFA kurtosis and electrode distances, *r* = −0.25, *p* < 0.001).

**Fig. 3.**
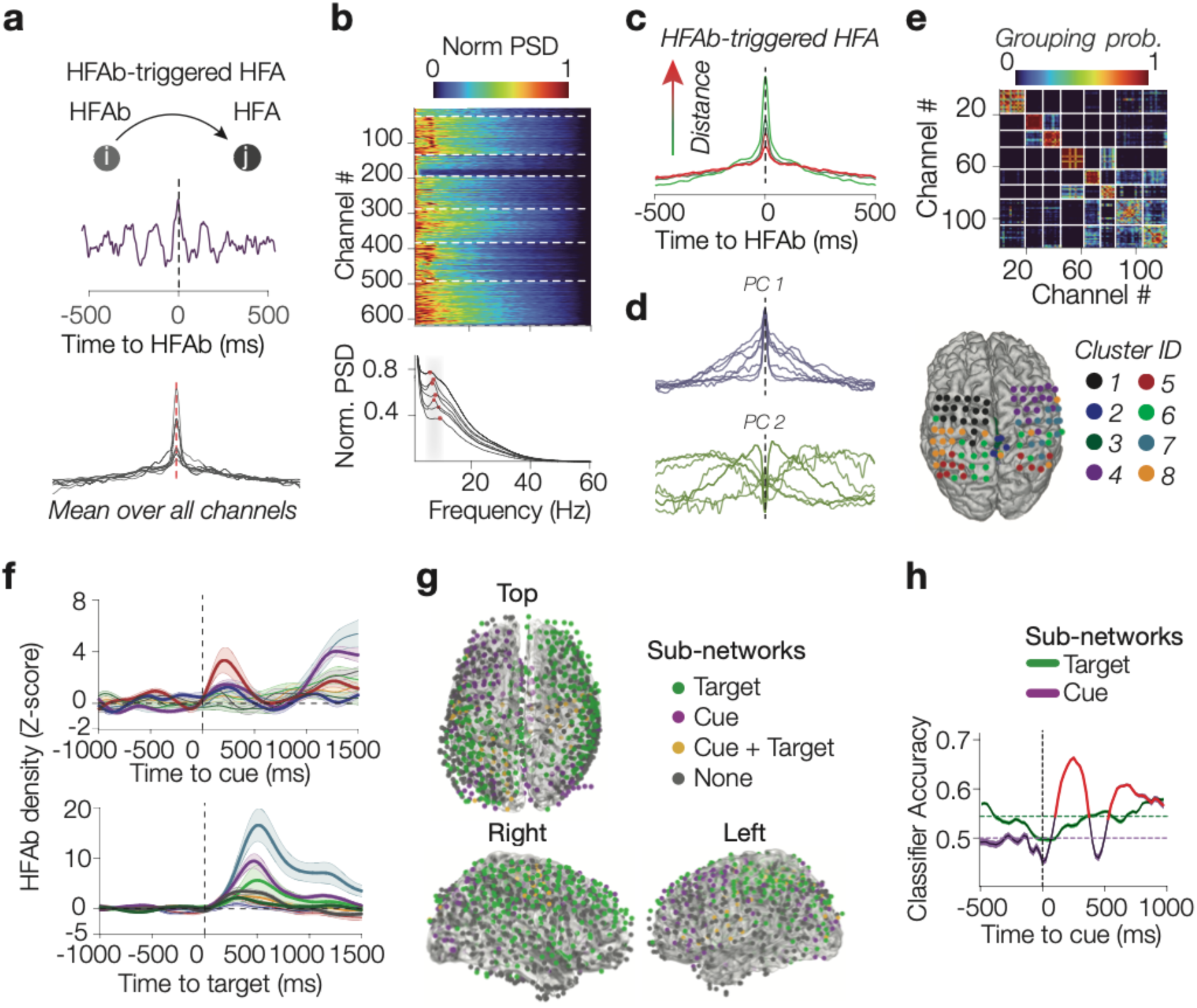
Network-level synchronization of HFAbs identifies functionally specialized subnetworks. **a**, HFAb-triggered HFA measured between electrode pairs during baseline periods (outside cue/delay and target/response epochs). Top: example electrode pair; bottom: group average across all pairs for individual subjects. **b**, Normalized power spectral density (PSD) of HFAb-triggered HFA for each electrode relative to all others (white dashed lines indicate subject boundaries), bottom: mean PSD across all electrodes for individual subjects (red dots denote peaks)**. c**, HFAb-triggered HFA HFAb-triggered HFA as a function of inter-electrode distance quartiles (25, 50, 75, and 100 mm), colored from green (short) to red (long). **d**, The first two principal components of HFAb-triggered HFA for individual subjects. **e**, Example subject showing pairwise grouping probability matrix for optimal K=8 clusters (top, white lines indicate cluster boundaries, clusters are ordered by stability, see **Methods**) and corresponding electrode topography (bottom). **f**, HFAb density around cue and target onsets for each cluster from panel **e** (mean ± SEM; thick lines indicate significant responses, *p* < 0.05, two-sided Wilcoxon test, FDR-corrected for multiple dependent comparisons). **g**, Group topography of electrodes, in clusters with significant cue responses ("cue-subnetworks") and target responses ("target-subnetworks") across all subjects. **h**, Performance classifier trained on mean HFAb density across electrodes in cue-subnetworks (purple) and target-subnetworks (green) around cue onset (error bars: SEM across realizations and cross-validations). Red lines indicate points significantly above baseline and chance (two-sided binomial test, *p* < 0.05, FDR-corrected for multiple dependent comparisons).

To extract temporal relations of high-frequency activity across electrode pairs, we used Principal Component Analysis (PCA) to reduce the dimensionality of HFAb-triggered HFA. The first component (PC1) reflected zero-lag synchronized and near-symmetric distributions of high-frequency activity in all subjects, explaining over 20% of the total variance (**Fig. 3d**, see **Extended Data Fig. 4d** for experiment 2). We projected HFAb-triggered HFAs onto PC space and used a resampling-based consensus K-means clustering technique to identify robust clusters of electrodes based on their scores on the synchronized PC (**Extended Data Fig. 4f**, see **Methods**). The clustering algorithm identified the most stable clusters (electrodes consistently grouped together across multiple runs) for each cluster number (K = 2-8, **Fig. 3e, Extended Data Fig. 5**). A series of accuracy metrics was used to determine the optimal number of clusters for each subject through a voting poll, selecting the cluster number that outperformed others across the majority of metric (**Extended Data Fig. 6a-c**).

Next, we examined whether these clusters were functionally specialized, revealing functional subnetworks within the larger brain networks. All subjects (except one excluded due to insufficient electrodes) showed clusters with distinct cue- and target-evoked activation profiles (**Fig. 3f**, see **Extended Data Fig. 6d** for individual examples). Clusters activated by cues and targets were labeled as *cue-* or *target-subnetworks*, respectively (Wilcoxon signed-rank test, *p* < 0.05; FDR corrected). Cue-subnetworks were predominantly located in occipital and parietal regions (e.g., IPS, TPJ, SPL, IPL), whereas target-subnetworks were more widely distributed across different brain areas including parietal, motor, premotor, and frontal cortices (**Fig. 3g**, see **Extended Data Table 1** for individual subject electrode locations).

Lastly, similar to **Fig. 1h**, we tested whether cue responses in cue- and target-subnetworks could predict trial outcomes. HFAb density averaged across electrodes in cue-subnetworks within 98-374 ms post-cue predicted successful detection of upcoming targets (binomial test, P < 0.001, FDR corrected for dependent samples). This prediction was specific to cue-subnetwork (**Fig. 3h**) and was not driven by a single subject (see **Extended Data Fig. 6e-g** for classifier details in both experiments). It is worth noting that HFAbs in both cue- and target-subnetworks, predicted trial outcome following target onset (**Extended Data Fig. 6h)**.

As a control, we re-referenced datasets from common average referencing to local composite referencing (LCR, each electrode was referenced to its nearest neighbors, see **Methods**), This re-referencing did not alter any of the main results. Together, these findings suggest that large-scale brain networks show synchronized HFAb patterns, with their synchronization identifying functionally specialized subnetworks that exhibit distinct temporal dynamics. Furthermore, HFAbs in cue-subnetworks following cue onset were predictive of performance accuracy.

### HFAbs in cue-subnetworks precede target-subnetworks

Our observation that HFAb responses in cue-subnetworks predict performance suggests their involvement in target detection, raising a key question: do cue-subnetworks route information to target-subnetworks during target processing, implying directional flow that supports attentional performance? To address this question, we used two different approaches (only subjects whose electrode coverage included both cue- and target-subnetworks were considered; n = 3 in experiment 1, and n = 3 in experiment 2). First, we measured HFAb-triggered HFA during target-to-response period and tested whether HFAbs showed temporally ordered activity between cue and target-subnetworks within this window. On average, HFAbs in cue-subnetworks preceded HFA in target-subnetworks during target processing (**Fig. 4a**, permutation test, *p* < 0.05, see **Extended Data Fig. 7a** for individual example). We further quantified the distribution of HFAb-triggered HFA peaks for individual subjects. The HFA peak distribution in target-subnetworks showed a mean at 39.7 ± 4.4 ms (n = 3) after HFAbs in cue-subnetworks and was significantly different from the opposite direction (**Fig. 4b**, Wilcoxon rank-sum test, *p* < 0.05). Similar results were found in experiment 2 (**Extended Data Fig. 7b**). For comparison, no lead-lag patterns were observed during the cue-to-target interval **(Extended Data Fig. 7c).** Topographically, and at the level of individual subjects, HFAbs in the occipital, posterior parietal, and frontal areas led over the motor/premotor areas during target processing (**Fig. 4c**, see **Extended Data Fig. 7d** for more examples).

**Fig. 4.**
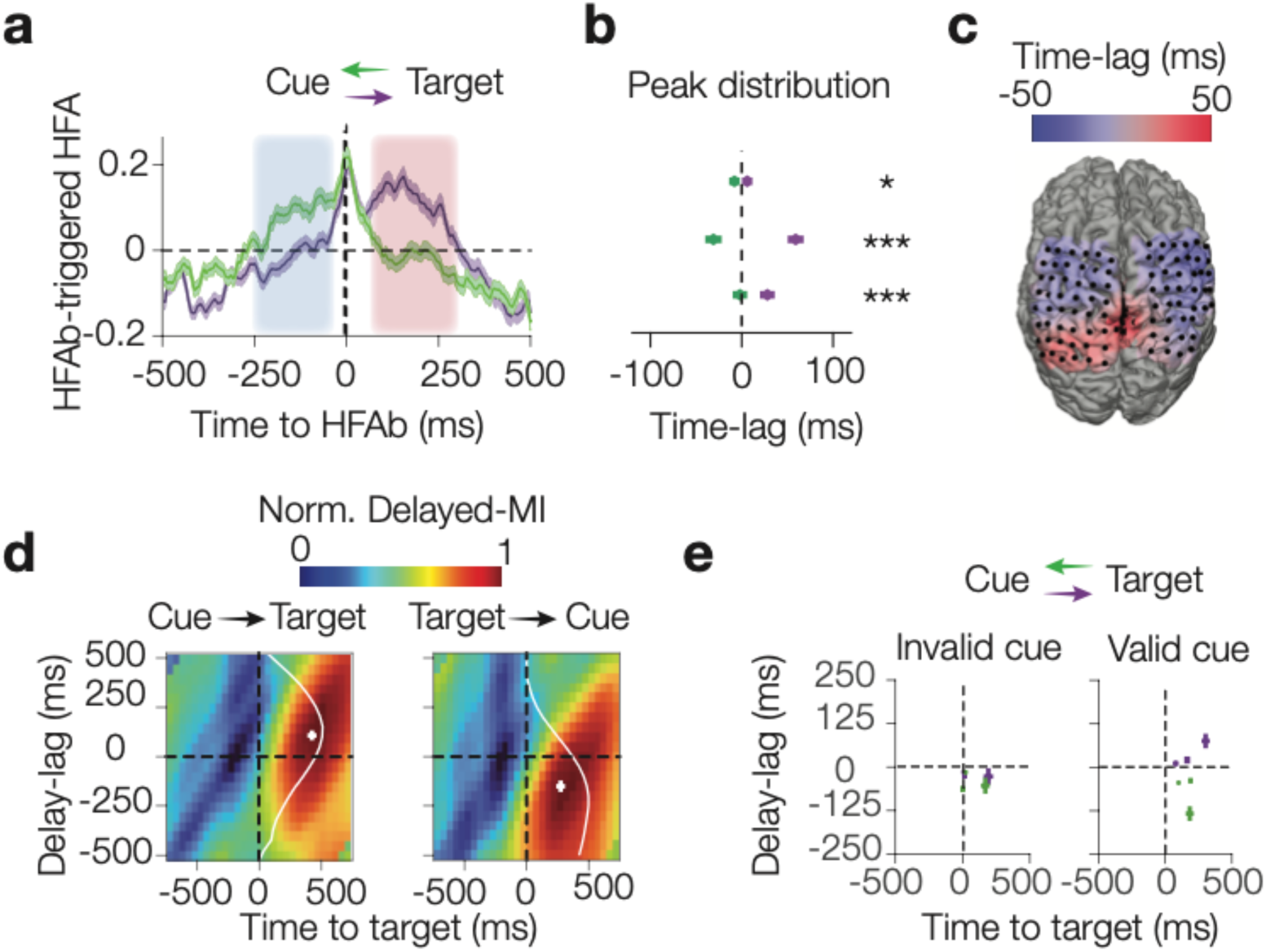
Cue-subnetworks precede target-subnetworks during target processing. **a,** Group-average HFAb-triggered HFA during target processing showing temporal relationships between subnetworks (mean ± SEM): HFA in target-subnetworks aligned to HFAbs in cue-subnetworks (purple) versus HFA in cue-subnetworks aligned to HFAbs in target-subnetworks (green). Shaded regions indicate significant lead-lag patterns (two-sided permutation test, *p* < 0.05; red: cue leads, [65 313] ms, blue: target leads, [-250 −37] ms). **b**, Time-lag distributions of HFAb-triggered HFA peaks for individual subjects (medina ± SEM; n=3, two-sided Wilcoxon rank-sum test between directions; *: *p* < 0.05, **: *p* < 0.01, ***: *p* < 0.001). **c**, An individual example of topographic visualization of lead-lag patterns between cue and target subnetwork electrodes averaged across cluster numbers during target processing period (see **Extended Data** Fig. 7 for more examples). The lead and lag temporal patterns are shown in a color gradient from red to blue. **d**, Group average heatmaps of normalized delayed mutual information (DMI) heatmaps showing temporal dependencies when shifting cue-subnetwork relative to fixed target-subnetwork (left) and vice versa (right). The white line indicates the time-lag distribution of DMI peaks across electrode pairs. The cross signs indicate the median of DMI peak time-lags ± SEM. **e**, Comparisons of DMI peak distributions between the two opposite directions (green for shifting the cue-subnetwork and purple for shifting the target-subnetwork) for valid (right) and invalid cue (left) trials. Individual subjects shown as crosses (n=3, median ± SEM across electrode pairs).

Next, we used delayed mutual information (DMI) to further quantify temporal dependencies between cue and target-subnetworks. DMI revealed (i) when cue and target-subnetworks shared maximum information relative to target onset, and (ii) time-lags of maximum inter-predictability. DMI peaked 285.4 ± 18 ms after target onset (**Fig. 4d**; n = 3). Time-lag distributions showed cue-subnetworks preceded target-subnetworks by 48.8 ± 23 ms during target processing (**Fig. 4d**, Wilcoxon signed-rank test, *p* < 0.001, n = 3; see **Extended Data Fig. 7e** for individual heatmaps). The temporal precedence occurred only after target onset, not during cue processing (**Extended Data Fig. 7f**), and specific to valid-cue trials, indicating it is not merely visual hierarchy activation, but reflects routing only when cue information is relevant for target detection (**Fig. 4e**). This pattern was consistent across individuals and both experiments (**Extended Data Fig. 7e-h**). Overall, these results demonstrate that subnetworks, identified by their network-level HFAb synchronization patterns show temporal sequences during task events. Specifically, when cues provide valid spatial information about the target, HFAbs in cue-subnetworks precede those in target-subnetworks following target onset.

### Computational modeling of HFAbs through spiking neural networks

We first developed a computational model using spiking neural networks to investigate both the mechanisms underlying HFAbs and their functional role in attentional performance. We simulated two interconnected networks, each consisting of 1000 neurons (80% excitatory, 20% inhibitory; **Fig. 5a, Extended Data Fig. 8a,** see **Methods**). Each network was fed by external input currents primarily to the excitatory population.

**Fig. 5.**
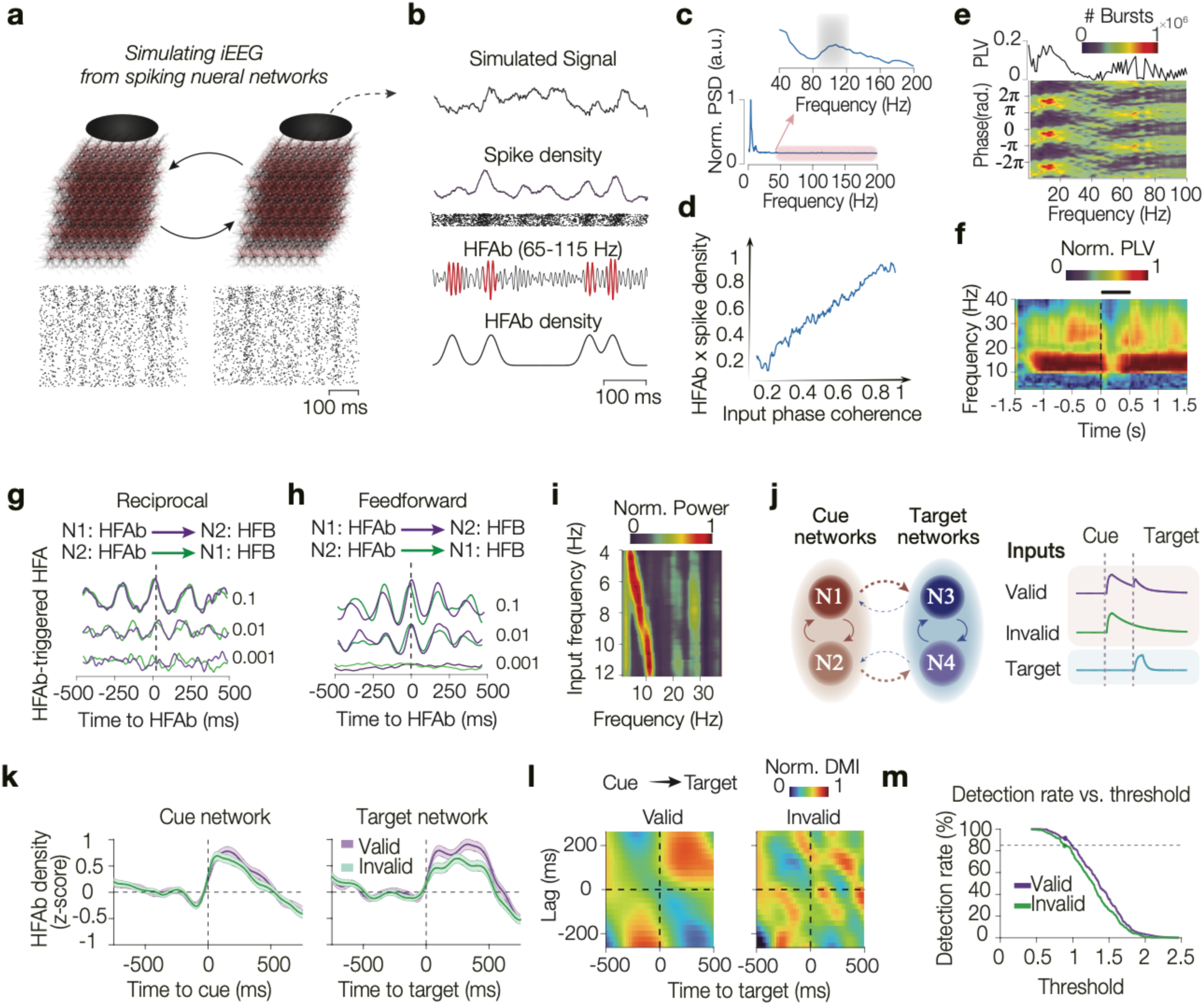
Computational modeling of HFAbs in spiking neural networks. **a**, Two interconnected cubic networks of point source 1000 spiking neurons 1000 (80% excitatory, 20% inhibitory) with recording electrodes measuring field potentials above each network (bottom: raster plots) **b**, Example trial showing simulated iEEG signal, population spike density with raster, detected HFAbs (red), and resulting HFAb density. **c**, The power spectral density (PSD) of iEEG signals. **d**, Correlation between HFAb and spike density increases linearly with external input coherence. **e**, HFAb phase distribution at different frequencies (bottom), and Phase-locking value (PLV) across all HFAbs (top). **f**, PLV of HFAbs to low frequencies following transient input (thick black line). **g,h**, HFAb-triggered HFA between reciprocally connected (**g**) and feedforward (**h**) networks with varying connectivity strengths (purple and green traces show HFAb-triggered HFA when HFAbs are extracted from N1 and N2, respectively). **i**, PSD over the HFAb-triggered HFA for rhythmic inputs of varying frequencies. **j,** Four-network model simulating spatial attention task: two spatially selective cue networks (N1, N2) with dominant feedforward projections to target networks (N3, N4); cue and target networks activate to cue and target onsets, respectively, cue networks reactivate when targets appear at cued locations. **k**, HFAb responses in cue networks to cue onset (left) and target networks to target onset (right) for valid (purple) and invalid (green) trials (mean ± SEM). **l,** Delayed mutual information heatmaps between cue and target networks for valid (left) and invalid (right) conditions showing temporal precision of cue over target networks following target onset. **m,** Detection rates as a function of HFAb threshold demonstrating enhanced detection for valid versus invalid trials.

We implemented two recording sites on top of each network measuring electrical field dynamics of postsynaptic and transmembrane potentials (**Fig. 5a,b**, **Extended Data Fig. 8b,c,** see **Methods**). Simulated iEEG signals showed low-frequency spectral peaks corresponding to network resonance frequencies and high-frequency (65–115 Hz) peaks, consistent with experimental data (**Fig. 5c**). To investigate the neural mechanisms underlying HFAbs, we calculated the spike density for the population of neurons, and the HFAb density from each recording site. HFAb density correlated with spike density compared to random timings (randomization test, *p* < 0.05) consistent with previous empirical findings.^25,28,38^ When feeding the network different levels of external non-rhythmic current coherence (**Extended Data Fig. 8c)**, we observed a linear increase in the correlation between HFAb and spike densities as input coherence increased (Spearman correlation, *r* = 0.87, *p* < 0.001, **Fig. 5d**). These findings suggest that HFAbs emerge when coherent external inputs drive neural populations into elevated firing states, indicating that HFAbs reflect input-driven population excitability rather than spontaneous fluctuations in neural activation.

Next, we examined HFAb phase-locking to slower rhythms. HFAbs triggered spectral peaks and showed phase locking to theta, alpha and beta bands (4-25 Hz, **Fig. 5e**), depending on synaptic time constants, external input strength, and neural connectivity strength, consistent with previous computational findings.^39^ To test network mechanisms underlying decoupling during cue and target processing, we modeled feedforward networks and measured time-resolved PLV in both networks after feeding one with a brief input pulse (∼50% of neurons receiving brief in-phase inputs; **Fig. 5f, Extended Data Fig. 8e,f**). The external impulse increased HFAb rate and desynchronized both networks, resulting in transient decoupling of HFAbs from low-frequency rhythms (Wilcoxon rank-sum test, *p* < 0.001, **Fig. 5i, Extended Data Fig. 8g,h**). These results reveal that external inputs shift neural networks from internally regulated states (HFAbs phase-locked to slow rhythms) to externally driven states, mirroring the empirical decoupling observed during cue and target processing.

We also examined HFAb synchronization patterns between networks. Reciprocally connected networks and networks with correlated external inputs showed synchronized high frequency activity (**Fig. 5g**), while feedforward networks showed lead-lag patterns within 150 ms of HFAbs (Wilcoxon rank-sum test, *p* < 0.001, **Fig. 5h**). Overall, these inter-network HFAb patterns showed intrinsically generated rhythmic fluctuations, though rhythmic external inputs could also entrain its rhythms to HFAb synchronization patterns between networks (Spearman correlation, *r* = 0.83, *p* < 0.001, **Fig. 5i**). These findings suggest that inter-network HFAb patterns reflect both structural connectivity and shared input statistics, providing mechanistic support for the zero-lag synchronization within subnetworks and lead-lag patterns between subnetworks observed empirically.

To link network dynamics to behavioral performance, we extended our model to a four-network architecture with spatially selective cue networks connected to target networks. All networks were activated by a shared decaying input initiated at trial onset. Cue networks responded transiently to cue onset and decayed thereafter. Target presentation was modeled as an additional input to the target networks. If targets appeared at cued locations, cue networks showed weak reactivation (**Fig. 5j, Extended Data Fig. 8i,j**, see **Methods**). We ran the model for 150 trials with valid and invalid cue conditions. Valid cues increased burst response in target networks following target onsets (Wilcoxon rank-sum test, p < 0.001, **Fig. 5k**). Delayed mutual information analysis revealed that cue networks temporally preceded target networks specifically for valid trials, with maximal information transfer occurring after target onset (**Fig. 5l, Extended Data Fig. 8k**), mirroring our empirical findings of cue-subnetwork precedence. We implemented a detection threshold model to test how cue-driven input enhances target detection. When fitting detection thresholds to invalid trials, valid cues consistently increased HFAb rates above threshold, resulting in higher detection probability (**Fig. 5m, Extended Data Fig. 8l**). This provides a mechanistic account of how cue information enhances behavioral performance through increased state transition probability by inputs from cue networks.

Our modeling results demonstrate that HFAbs reflect transitions in neural populations into spiking states. HFAbs are phase locked to low-frequency rhythms, but transient inputs disrupt this coupling. HFAb synchronization patterns between networks depend on connectivity and task structure, with behavioral relevance demonstrated through enhanced detection rates when cue-driven inputs increase target network burst rates.

## Discussion

Here we present a framework for understanding mechanisms of large-scale information routing in brain networks at fast timescales. We characterize high-frequency activities as discrete burst events (HFAbs), rather than continuous fluctuations, supported by computational and animal electrophysiology studies emphasizing their circuit-level origins and roles in long-range communication.^6,40–44^ This quantification allows a millisecond-resolution, network-level account of selective attention that extends beyond traditional pairwise interareal interactions. Our findings revealed that HFAbs were synchronized across distributed cortical networks, identifying functionally specialized subnetworks for cue and target processing. Importantly, HFAbs in cue-subnetworks following sensory cues predicted trial-by-trial performance accuracy and temporally preceded HFAbs in target-subnetworks during target processing when cues were informative. Complementing these findings, our computational modeling showed HFAbs emerge from bouts of elevated excitatory drive to local networks, marking fast state transitions in neural populations that support attention processes.

Our analysis of both exogenous and endogenous attention tasks revealed conserved mechanisms despite distinct control pathways. In both tasks, valid cue information enhanced target-triggered HFAb responses, and cue subnetworks showed temporal precedence over target subnetworks only when the cue was informative. These preserved findings across both tasks suggest that information routing signified by HFAb dynamics represents a fundamental mechanism for transforming cue information into spatial priority signals, regardless of whether attention is stimulus-driven or top-down controlled.^45,46^ However, task-specific differences emerged in the timing of lateralized processing; exogenous attention showed laterality effects during cue processing, reflecting stimulus-driven nature of peripheral cues, while endogenous attention showed laterality only during target processing, consistent with the internal maintenance and deployment of spatial attention from symbolic cues.^46^ These findings support the argument that while attentional control signals may rely on different cortical pathways, more posterior for exogenous and involving frontal regions for endogenous attention, they converge onto common selection mechanisms in parietal and sensory areas.^47,48^

A main finding of our study revealed that the most prominent pattern of high-frequency dynamics in the brain network was zero-lag synchronization (**Fig. 3d**), consistent with previous studies showing phase locking of high-frequency dynamics in brain networks.^31–33^ Zero-lag synchronization patterns are thought to enhance large-scale information processing and facilitate representational states of sensory information in the brain.^49–51^ While our results cannot directly identify the origins of these synchronized interactions, two mechanisms might explain these patterns: subcortical inputs,^52–55^ particularly higher-order thalamic nuclei,^52,56–58^ or direct intracortical interactions.^29,59–61^ Our modeling supports both these possibilities showing correlated inputs to distinct neural networks, or reciprocal connections between spiking networks could drive such long-range synchronization (**Fig. 5g**).

We further showed that network-level baseline synchronization of high-frequency dynamics reliably identifies distinct functional brain subnetworks. While the baseline state did not contain cue or target processing periods, it could identify subnetworks with unique activation patterns in response to those behavioral events (**Fig. 3**). This approach diverges from conventional methods for delineating brain network organizations and communications, such as fMRI, which primarily focuses on low-frequency BOLD signal fluctuations.^62,63^ By leveraging high-frequency dynamics, we demonstrate a link between baseline neural activity and task-induced network activation patterns, suggesting that intrinsic network fluctuations can affect behavioral responses at millisecond timescales. Identifying these subnetworks at fast time scales is critical to capture rapid state transitions in brain networks, which may underlie the temporal structures necessary for adaptive information routing during behaviorally relevant events.^64^ Our findings demonstrate that brain subnetworks, identified by their network-level HFAb synchronization, exhibit lead-lag patterns during behavioral events. Specifically, cue-subnetworks preceded target-subnetworks during target presentation when spatial cues provided relevant information about target locations (**Fig. 4**). Our modeling provided a mechanistic account for this, showing that inputs from cue networks during target processing increase the probability of state transitions in target networks, reflected by elevated HFAb rates (**Fig. 5k,m**). Overall, these observations uniquely bridge the gap between studies focusing on local and cross-regional mechanisms of fast information routing^6,36,41,42,44,56^ and those examining network dynamics beyond pairwise interactions,^15,65^ providing new insight into the mechanisms of selective attention.

Another key observation was the coupling dynamics of HFAbs and local low-frequency activities. We found that phase-locking of HFAbs to low-frequency dynamics occurs even with random external input patterns in modeling, and this is a widespread phenomenon in iEEG signals, consistent with previous studies.^30,66,67^ However, our findings further showed that HFAbs transiently decouple from local low-frequency phases in response to both cue and target events (**Fig. 2**). Our modeling supports this observation, showing that transient inputs interrupt the default state coupling of HFAbs to low frequencies (**Fig. 5f**). These network interferences, characterized by brief and strong responses, can temporarily induce local cross-frequency decoupling, possibly by increasing response heterogeneity and disrupting the timing of inhibitory and excitatory neuron activation.^68^ This desynchronization may enhance cognitive processes such as selective attention, perception, and memory retrieval by facilitating the processing of new information and suppressing internally regulated activity states.^68–71^ We observed that this decoupling, accompanied by increased HFAb density following cue and target events, resembled a desynchronized up-state, a brain state characterized by activated but desynchronized neural activity across cortical networks. Importantly, this decoupling was more pronounced in correct trials during both cue and target processing, suggesting that desynchronization correlates with accurate cognitive performance (**Fig. 2f**).

It is worth noting that the HFAbs identified in this study share spectral features with cortical ripples recently described in human studies,^33,38,72^ However, HFAbs exhibit different characteristics suggesting specialized functions. While both show synchronization and correlate with population spiking, HFAbs in our data occurred ∼15-fold more frequently than ripples, with shorter durations (∼36 vs 50-100 ms).^33,72^ Ripples are largely described in support of memory maintenance with consistent phase-locking to slow waves,^72,73^ whereas HFAbs show performance-dependent decoupling from oscillations and directional lead-lag patterns between functional subnetworks. These distinctions suggest that ripple-band high-frequency bursts constitute a general population-level mechanism that can flexibly switch between binding (memory) and routing (attention) modes, with HFAbs instantiating the routing mode during active sensory processing.

In summary, our study provides a multilevel approach to understanding large-scale cortical communications, showing that HFAbs are markers of fast population state transitions that enable selective information routing through zero-lag synchronization within subnetworks and lead-lag temporal sequences between subnetworks.

## Acknowledgments

This work was supported by the C.V. Starr Fellowship (KBB), German Research Foundation, Emmy Noether Program (DFG HE8329/2-1; RFH), NIBIB (P41-EB018783; PB), NINDS (R01NS021135; RTK), and NEI (2R01EY017699; SK), NIH (1R01MH137624; SK), NIMH (2R01MH064043, P50MH132642; SK). The funders had no role in study design, data collection and analysis, the decision to publish, or the preparation of this manuscript.

## Author Contributions

Conceptualization, K.B.B. and S.K.; Methodology, K.B.B.; Formal Analysis, K.B.B.; Modeling, K.B.B.; Investigation, R.F.H., R.T.K. and J.J.L.; Software, R.F.H. and I.C.F.; Visualization, K.B.B; Writing – Original Draft, K.B.B.; Writing – Review & Editing, K.B.B., R.F.H., I.C.F., J.N.B., P.B., J.J.L., R.T.K., and S.K..; Supervision, S.K.

## Methods

### Experimental model and subject details

#### Participants

Intracranial recordings were obtained from 12 epilepsy patients who underwent pre-surgical monitoring with implanted grid electrodes. Study 1 included seven patients (35.99 ± 12.42 years; mean ± SD; 5 females; see^23^ for further details). Patients were recruited from the University of California, Irvine Medical Center, USA (n = 6) and California Pacific Medical Center (CPMC), San Francisco, USA (n = 1). Study 2 included 5 patients (30.20 ± 1186 years; mean ± SD; 1 female, 3 patients were excluded from the original study due to their limited electrode coverages; see^3^ for further details) from Johns Hopkins Hospital in Baltimore, MD, USA (N = 1) and Stanford Hospital, CA, USA (N = 4). The electrode placement was entirely guided by clinical considerations, and all patients provided written informed consent to participate in the study. All procedures were approved by the Institutional Review Board at each site, as well as the Committee for Protection of Human Subjects at the University of California, Berkeley (Protocol number: 2010-02-783) and were in accordance with the Declaration of Helsinki.

### Experimental design and procedure

#### Behavioral tasks

Participants performed a spatial attention task in each experiment. In experiment 1, participants performed a variant of the Egly-Driver task;^74,75^ see^23^ for further details) Behavioral data were collected with Presentation Software (Neurobehavioral Systems). Subjects were seated approximately 60 cm from the laptop screen. Subjects started each trial by pressing a left mouse button. On each trial, after appearance of a fixation cross, two bars appeared vertically or horizontally on the screen, followed by a brief spatial cue (100 ms), presented after 400-800 ms at one of the four corners of the bar stimuli. The spatial cue indicated the location where the target was most likely to appear (72% cue validity) and occurred pseudo-randomly in any of the four quadrants. A variable cue-to-target interval (500 – 1700 ms) was introduced after the cue during which participants sustained spatial attention at the cued location. Targets could randomly appear at any point during the cue-target interval, and participants released the mouse button to report a detected target. Infrequent catch trials (10%) during which no actual target appeared were used to track the false alarm rate. Auditory feedback indicated whether the trial was performed correctly. The target luminance was adjusted every 15 trials, if necessary, by increasing/decreasing the RGB value, in order to achieve an overall approximate accuracy of 80%. The experimenter monitored continuous fixation. All participants responded by using the hand contralateral to the implanted grid, except for participant S5 who had bilateral grids and responded by using the left hand. Participants performed up to 5 blocks of 60 trials each (190 trials ± 67; mean ± SD).

Experiment 2 (Extended Data Fig. 1) used EPrime software (Psychology Software Tools) to control stimulus presentation (see Szczepanski et al., 2014 for more details). Subjects were seated approximately 60 cm away from the laptop screen. Each trial began with red circles (distractors) dynamically switching on and off on a dark background. A spatial cue guided participants to the right or left hemifield, and the cue remained on the screen throughout the trial. Subjects were instructed to maintain fixation and only covertly shift their attention to the cued hemifield. Through the trial, the experimenter monitored eye movements and ensured central fixation. After a variable cue-target interval (1000 – 2000 ms), a blue square target appeared on the screen (∼62/38% on the cued/uncued hemifields). The target remained on the screen until the subject responded or the trial ended (2000 ms timed out). Subjects were asked to report a target seen in a cued hemifield by pressing a button while withholding a response if the target was seen in a non-cued hemifield. Targets appeared randomly during the cue-target interval. Three out of five subjects responded with the hand ipsilateral to the grid. Each participant completed six blocks (each 42 trials). The experimenter monitored eye movements and ensured central fixation throughout both experiments.

#### ECoG data acquisition

Electrophysiological and peripheral (photodiode) data were collected using a Nihon Kohden recording system at UC Irvine, CPMC and Children’s Hospital (128/256 channel, 1000/5000 Hz sampling rate), a Tucker Davis Technologies recording system at Stanford (128 channel, 3052 Hz sampling rate), or a Natus Medical Stellate Harmonie recording system at Johns Hopkins (128 channel, 1000 Hz sampling rate).

#### Electrode localization

In experiment 1, the electrodes were localized by transforming both the pre-implant MRI and the post-implant computed tomography CT into Talairach space. For all subjects, MNI coordinates were also calculated for each electrode location, which was used for group-level visualizations.^23^ In experiment 2, post-implant CT was aligned to the pre-implant MRI and all were transformed into MNI space across subjects.^3^ Strip or grid electrodes were implanted with 1 cm spacing. One participant (S5) had an additional 8 contact depth probe inserted into the occipital cortex. Electrode positions were primarily determined using the VTPM atlas.^76^ For electrodes without an assigned label, the process was repeated using the AFNI atlas.^77^ The assigned positions were manually verified and adjusted based on electrode reconstructions visualized in native Talairach space. Electrodes near the Temporoparietal Junction (TPJ) were manually localized, as TPJ definitions were not available in either the VTPM or AFNI atlases (see Helfrich et al., 2018 for more details).

#### iEEG Data

Preprocessing: All intracranial EEG channels were manually examined by a neurologist for epileptiform activity and artifacts. Affected channels and epochs were excluded. The raw data was preprocessed using the EEGLAB and Fieldtrip toolbox Field Trip toolbox^78^ Version 20230118 in MATLAB, The Mathworks Inc. R2023b; https://www.mathworks.com/.

Preprocessing included notch filtering at 60 Hz and all harmonics, as well as referencing the data to the common mean of electrodes.^3,23^ Then, the data was time locked to individual trials. Trials were 8 seconds long, −3 to +5 seconds around cue onsets in the experiment 1 and −2 to +6 seconds around cue onsets in the experiment 2. As a control, we re-referenced the datasets from common average referencing to local composite referencing (LCR), a spatial Laplacian estimate relative to nearest neighbors. This re-referencing did not alter any of the main results.

### Analysis of high frequency activity bursts (HFAbs)

#### HFAb detection and density analysis

We adopted an adaptive burst detection approach, similar to previous work,^6^ to identify high-frequency oscillatory bursts (65-115 Hz) at each electrode. The frequency band was selected based on spectral peaks observed across all individuals (87 Hz trials ± 14; mean ± SD) and in the modeling. Similar analyses were conducted for a broader band (65-175 Hz) and the low gamma band (35-65 Hz). While the broader band produced similar results, the low gamma band yielded inconsistent responses and noisy clustering. We selected the frequency band based on spectral peaks across individuals and in the modeling. First, we applied a zero-phase Butterworth bandpass filter to the padded signal and then calculated the analytic signal x(t) using the Hilbert transform. We extracted the instantaneous amplitude as the real part of the analytic signal *z*(*t*) following eq. 1:

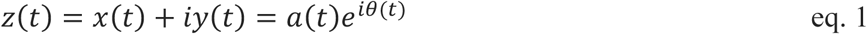

where *y*(*t*) is the Hilbert transform of *x*(*t*):

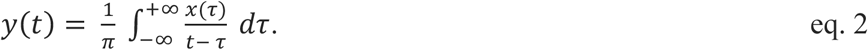

The instantaneous energy ^79^ *IE*(*t*) of the signal is calculated from its Hilbert transform:

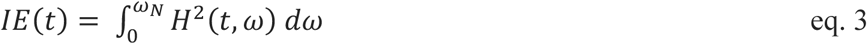

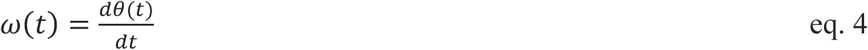

where *ω*(*t*) corresponds to instantaneous angular velocity. If the frequency band is narrow and if the instantaneous frequency (eq. 4) is small enough, it can be approximated by squared *a*(*t*), which is the instantaneous amplitude. The marginal mean energy of the signal then can be estimated as

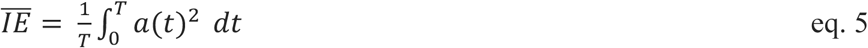

where T represents the duration of the signal. For skewed energy distributions in empirical data, we determined a median absolute deviation (MAD) threshold by setting it to 3.3 times plus the median energy.

To qualify as a burst, the energy level had to exceed this threshold for at least 1.5 cycles of the upper bond frequency, and the instantaneous amplitude needed to surpass 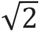 times the RMS of the peaks. Burst boundaries were marked by the closest points to a burst peak where either the instantaneous energy fell below the signal’s mean energy or the deviation in instantaneous frequency exceeded the mean change plus two standard deviations. This was to exclude multi-component or noisy events. Bursts were considered significant if their duration was at least 2.5 times the upper bond frequency cycle and exceeded the average span of adjacent local minima of the energy function. Finally, when bursts were too close (less than five frequency cycles apart), only the burst with the higher energy peak was kept. For comparison to ripple, we also quantified ripples in our data by calculating z-score of instantaneous energy, and finding events lasting longer than 3 cycles above a value of 2.5.^33^ We observed extremely rare events in our data with ripple characteristics.

To estimate how HFAb events are distributed over time, we calculate the HFAb density by convolving a vector of burst events with a gaussian window of 500 ms and a standard deviation of 100ms. After estimating the HFAb density for each channel and trial, we investigated whether the cue and target stimuli affected burst density. For each electrode, the average burst density aligned to the cue and target onsets were calculated separately. Following that, we normalized timeseries to the baseline (1000 ms before each event onset) by subtracting the mean and dividing it by the standard deviation (see **Fig. 1c**). This is a similar approach to the peri-stimulus time histogram (PSTH) for spike trains ^80^.

#### Visualization of individual and group average responses on 3D brains

To visualize how HFAb responses were topographically organized in the 3D brains, we plotted both individual (**Extended Data Fig. 1f**) and group responses to cue (**Fig. 1f**) and target onset (**Fig. 1g**). Each electrode value was calculated as the mean of Z-scores within 500 ms of cue and target onset. For individual subjects, the plotted value for each electrode was linearly attenuated with distance from the electrode and for a sphere of 1 cm radius (illustrating correct versus incorrect trials separately, **Fig. 1h**, **Extended Data Fig. 1f**). The group average plot (**Fig. 1f,g, Extended Data Fig. 2e**) was calculated using electrode locations from both experiments to better cover the entire brain. The MNI coordinates of electrodes were used for rendering on a template brain (subject S4 from experiment 1 and subject S12 from experiment 2 were excluded due to suboptimal wrapping of Tal to MNI spaces). We set the value of all mesh surfaces to zero for each subject. Then, similar to the plotting for individual data, we calculated the value for mesh surfaces by using linear attenuation (the sphere radius was set to 2.5 cm in order to achieve a smoother visualization of the whole brain). As a last step, we averaged the surface values across all subjects to find consistent patterns of activations for the cue and target. We plotted the data onto 3D brain for correct and incorrect trials separately.

#### Statistical analysis of HFAb responses to cues and targets events

After calculating the burst density and finding PSTH for each channel, we examined whether the HFAb responses evoked by cues (within a window of 500 ms after cue onset) and targets (750 ms after target) were significantly different from baseline. This analysis was done at the network level. The average responses of electrodes within the defined time window were determined using the normalized HFAb density for each channel. Wilcoxon signed-rank tests were then used to examine if there was a non-zero response at the network level to cue and target onsets. Also, we performed a linear regression analysis of the cue and target responses in electrodes under the null hypothesis that the cue and target responses are independent (**Fig. 1e**).

To test whether cue and target responses were different when grouped by trial outcomes, we used the Kruskial Wallis test under the null hypothesis that cue, and target responses do not differ by outcome condition. For pairwise comparisons between different groups, we used Dunn’s test ^81^ with Tukey-Kramer multiple comparison correction (**Fig. 1d**).

#### Identifying cue-responsive electrodes

We identified Cue+/- electrodes by determining increased HFAb rate or density profile within window of 500 ms after cue onsets. For each electrode, we compared the average HFAb density following cue onset with that at baseline using Wilcoxon rank-sum test. Electrodes with significant increases in HFAb rate after cue onset were labeled as Cue+ electrodes, while the remaining electrodes were labeled as Cue-electrodes, which was subsequently used in the classification (**Fig. 1k**).

#### Generalized Linear Mixed Effect Models

We used generalized linear mixed effects (GLMEs) models to examine the effects of each task variable on HFAb response to cue and target events. Three main predictors were used for the independent variables: *Laterality* (if the cue was either ipsilateral or contralateral to the electrode) with two levels (*Ipsi* and *Contra*), *Validity* (if the target appeared at the cued location) with two levels (*valid* and *invalid*), and trial outcome with four levels (*Correct hit, False alarm, Correct Rejection, Miss*). Since not all subjects had trials for all four outcome conditions, an alternative analysis also considered outcome as a binary variable (*Correct* and *Incorrect*) with a logit link function. The random effect was determined by considering groups as subject and channel labels. The response variable was either the mean burst density in response to the target or the cue. The GLME is then formalized as shown in eq. 6:

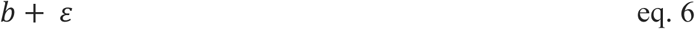

#### Training classifiers for predicting trial outcome based on HFAb density around cue onset

A time-resolved classification approach was used to determine how accurately HFAb responses evoked by sensory cues could predict trial outcomes. First, we selected electrodes that showed a significant HFAb responses to the cue presentation (*Cue+* electrodes, see above), for which we then calculated the average HFAb density for *Cue+* electrodes for each trial and subject. Next, we tested whether the average burst density in sliding windows around the cue onset could predict the outcome at the trial level with a sliding window of 350 ms and a step of 25 ms. We used binary Support Vector Machine (SVM) with one-to-one comparisons of HFAb density in sliding windows with five folds of cross-validation. For training each SVM, a vector of the HFAb density in each sliding window was used along with a vector of outcome labels (*"Correct"* = 1, *"Incorrect"* = 0). The classifier used a Gaussian radial basis function kernel with a scaling factor of one. We assured that the number of correct and incorrect trials used in the training was identical, to prevent sample size biases. As the number of incorrect trials were lower than the number of correct trials, we randomly sampled correct trials with a sampling size equal to the number of incorrect trials in each fold. For each sliding window, we trained the classifier 1000 times and calculated the average accuracy and confusion matrix across all iterations and folds. (**Fig. 1k).**

To determine whether HFAb density after cue onset could more accurately predict trial outcome than chance and baseline, we used a binomial test on the accuracy of each window against the maximum value of baseline and chance-level prediction accuracy. After obtaining a p-value for each binomial test, we corrected the p-values using false discovery rate (FDR) for dependent samples with a 0.05 alpha level ^82^ to correct for multiple comparisons error rate across all windows. We performed this procedure separately for each experiment (**Fig. 1k**). In our main analysis, we pooled data across all subjects in each experiment due to the low number of trials per subject. However, to ensure that the results were not solely dependent on one subject, we ran a control analysis in which one subject was left out each time and the classifier was trained and tested. Controlled analyses in both experiments demonstrated that our observation was not based solely on one subject. We used a similar approach for training the classifier on cue and target-subnetworks to train the classifier on cue and target-subnetworks, as further detailed in the ‘Identifying synchronous subnetworks’ section.

#### HFAb-triggered spectrum and Phase Locking Value (PLV) analysis

To understand the spectral dynamics of HFAbs and local population activities, we performed HFAb-triggered spectrum analyses using HFAb centers as discrete points. We estimated the PLV of points (burst centers) to their local field activity dynamics (both in the modeling and electrophysiological data) in order to investigate the phase synchronization of HFAbs within their local networks. We calculated the HFAb-triggered spectrum using an adaptive window around HFAbs. We extracted a window centered around each selected point (HFAb center), which covered 2.5 cycles of the frequency of interest before and after the selected point. This window was then multiplied by a Hanning window. We estimated the power spectrum for each window using the Fast Fourier Transform (FFT) for frequencies ranging from 1 Hz to 100 Hz. To account for trial-wise power variations, triggered spectrum estimates were normalized by dividing by total power. We averaged spectrum estimates across all points and trials to calculate the final HFAb-triggered spectrum. Following the analysis of the HFAb-triggered spectrum, a PLV calculation was performed to quantify the level of phase consistency of the HFAbs across trials for the frequencies of interest. The phase angle for each frequency was calculated based on the FFT results from the HFAb-triggered spectrum analysis. The PLV of HFAbs at each frequency was then calculated as the mean resultant in eq. 7:

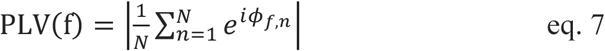

where N is the total number of HFAbs and *ϕ*_*f*,*n*_ denotes the phase angle for the n^th^ HFAb at frequency *f*. The statistical significance of the observed PLV values was determined using a non-parametric permutation test with 1000 permutations and the Rayleigh test corrected for multiple comparisons, at 0.05. Peaks that passed both tests and had a prominence of 25% higher than the PLV range were considered significant (shown in black dots in **Fig. 2c**). We further quantified the phases at peak frequencies (**Extended Data Fig. 2a**), peak width (**Extended Data Fig. 2d**), and the power ratio of peak frequency relative to baseline (**Extended Data Fig. 2e,f**) to ensure that HFAb phase-locking occurred within narrow frequency bands.

#### Time-resolved PLV analysis

To quantify the temporal dynamics of the phase synchronization of HFAbs with the low frequency LFP during cue and target processing, we extracted the phase of HFAbs at each frequency as explained above. Then, we used a sliding window of 500 ms with a step of 25 ms between −1500 and 1500 ms around the cue and target onsets separately. PLV was calculated for each sliding window and across all trials (different types of trials were analyzed separately, e.g., correct trials and incorrect trials). We set a minimum number of data points of 50 bursts for the analysis (to achieve reliable statistics on circular data,). We measured the PLV for 1000 samples of 25 HFAbs drawn randomly from a population of HFAbs in each time bin to control for the number of bursts. The PLV for each time bin was calculated as the average PLV across all randomly drawn samples. The average PLV value for each subject and for all subjects in each experiment was calculated separately.

To test whether HFAb phase synchronization differed during cue and target processing, we used a channel-specific randomization test. For each recording channel and time bin, 1000 subsamples of HFAbs were randomly selected from the baseline period (within 1 second before each event). For each channel, each time bin, and each frequency, we calculated the upper and lower bond CIs (2.5% most extreme PLVs). For multiple comparison correction, we repeated this procedure 1000 time and found the most extreme 2.5% value across all CIs for time bins under the null hypothesis that the PLV for each time bin and frequency is not significantly different from the baseline value. Each time-frequency bin was considered significant if it differed from the critical values (**Extended Data Fig. 3c**). Each electrode and frequency band (theta/alpha (4-14 Hz) and beta (15-25 Hz)) for a time bin was considered significant if it showed a significantly different PLV from the baseline in more than 25% of the frequency points in that frequency band. Next, we computed the mean number of electrodes with significantly different PLVs than the randomized PLV distribution for each subject. Using the binomial test, we determined whether the proportion of electrodes with significantly lower PLV after the cue and target onset was different from the baseline level as well as the chance level (5%, **Fig. 2d**). To adjust for multiple comparisons, we used FDR for dependent samples and an alpha level of 0.05.

We also performed a control analysis to ensure that event-evoked iEEG signals are not confounding the variation in synchronization. First, we extracted −1.5 to 1.5 seconds around cue- and target-aligned iEEG signals for each electrode and trial. We averaged the data across all trials and removed the average event-triggered iEEG signal from individual trials. The same synchronization and statistical analyses were then performed on the trial with event-evoked iEEG subtracted. Subtraction of event-evoked iEEG did not change the main results pattern (**Extended Data Fig. 3d**). While our main analyses controlled for HFAb counts, we also conducted a control analysis using pairwise phase consistency (PPC), a measure commonly used to quantify spike-field synchronization that is unbiased by event count. Results were consistent with our PLV analysis (**Extended Data Fig. 3e)**.^83^

#### Analysis of Coupling ratio

A Coupling Ratio (CR) index was defined to compare PLV after cue and target events as compared to baseline. We used this CR index also to visualize coupling variation in both 3D brain renderings (**Fig. 2e**), and the average network level coupling analysis (**Fig. 2g,h**). We calculated the coupling ratio by:

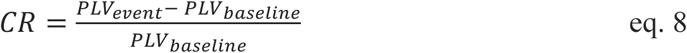

Where the *PLV*_*baseline*_ is the average PLV for each electrode within 1 second before the event, and *PLV*_*event*_ is the average PLV for each electrode within 0.5 seconds after the event. In this context, network decoupling is defined as a negative CR value indicating a reduction in PLV relative to the baseline. We show the CR value at both cue and target events for each subject.

We also used GLME models to investigate the effect of cue and target response on the coupling ratio following cue and target onsets (**Fig. 2g,h**). The GLME is formalized as shown in eq. 9:

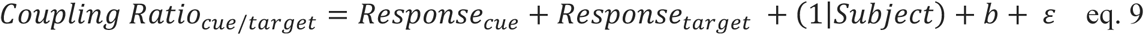

### Quantification and Spectral analysis of HFAb-triggered HFA

We calculated the interaction between HFAbs and HFA between each pair of electrodes. To estimate HFA, first we used a zero-phase Butterworth filter with cut-off frequencies of 65 Hz and 175 Hz to bandpass filter the signal. We chose the HFA frequency band broader to capture larger spectral content (similar results were obtained by choosing 65-115 Hz band). Next, we performed a Hilbert transform and used the real part of the analytical signal as HFA amplitude (eq. 1). We extracted a duration of one second around each HFAb from the HFA signal. We then calculated the HFAb-triggered HFA for each individual burst event as well as the average value for all bursts between each pair of electrodes. Our main analyses were restricted to HFAb-triggered HFA that occurred outside of the main behavioral epochs; either before cues or after target detection (**Extended Data Fig. 4e**, for those without a response 1 second following target presentation). To control for any burst-independent correlations of iEEG signals across electrodes, we first calculated the HFAb-triggered HFA using randomly assigned burst times. Each burst was jittered with a random time lag of ±1000 ms. We then subtracted this jittered HFAb-triggered HFA from the original for further analysis.

We calculated the spectral power of HFAb-triggered HFA across all electrode pairs to assess how HFA is organized relative to HFAbs recorded on other channels. The power spectral density was computed over the ±500 ms time window using a hanning taper. Additionally, we analyzed peak prominence for all pairs and plotted the distribution of spectral peaks. For each pair of electrodes, we extracted the prominent spectral peaks and plotted their distribution.

#### Dimensionality reduction of high-frequency temporal relation patterns

To identify prominent temporal patterns of high-frequency activities within the brain network, we analyzed a high-dimensional space of HFAb-triggered HFA across all electrode pairs for each subject, with each electrode pair representing a single dimension. Using the HFAb-triggered HFA population, we calculated the covariance matrix C by:

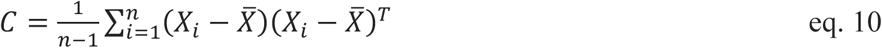

where *X*_*i*_ shows each HFAb-triggered HFA time series, n is the total number of electrode pairs, *X̄* is the marginal mean, and *X*^*T*^ is the transpose matrix of *X*. We found the eigenvectors of C as in eq. 11:

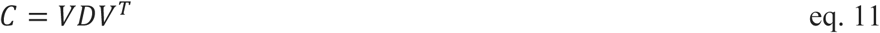

where V columns are the eigenvectors and principal components (PCs), and D contains eigenvalues indicating how much variance each component explains (**Fig. 3d**). In all participants, PC1 showed a near-symmetric activity state and explained more than 10% of variance in HFAb-triggered HFA dynamics over the population of electrode pairs (we used multistep Wilcoxon rank-sum test to examine if there was asymmetry of HFA within mirrored time windows around the HFAbs). By projecting the original data to the PC vector space, we calculated the HFAb-triggered HFA scores as:

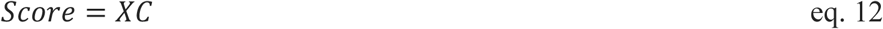

We then used the scores of electrode pairs on the first component as their loading values on the synchronous component of the population (**Extended Data Fig. 4**).

#### Identifying synchronous subnetworks

Based on the scores of each electrode pair on the synchronous principal component, we generated a network synchrony matrix that describes synchronous inter-electrode interactions. For each electrode, we then defined a vector of variables with a dimension equal to the total number of electrodes. Each element of this vector describes the score of an electrode pair on the synchronous component. This vector was defined for all electrodes, which generated a matrix in which each row represented one observation (electrode) and each column represented the score of the observed electrode on another electrode.

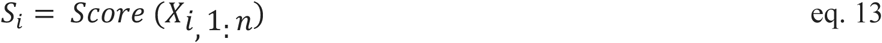

Using this matrix, we clustered the electrodes using resampling-based consensus K-Means algorithm with a correlational defined distance as in eq. 14:

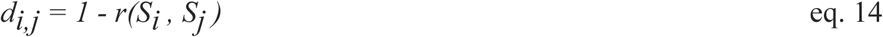

where *d_i,j_* is the correlation distance between two electrodes *i* and *j*. The correlational distance between two electrodes is low when they show synchronous activity, and similar synchronization patterns to the rest of the network. First, we used several clustering indices, including Hubert, Silhouette, Davies-Bouldin, Calinski-Harabasz, Hartigan, Homogeneity, and Gap, to find an optimal range of cluster numbers (between 2 and 8 for all subjects) ^84–87^.

For each cluster number, we ran the K-means algorithm 1000 times. The sample size was subsampled and only 25 percent of electrodes were randomly selected for each clustering (we ensured that each cluster had at least an average of 5 data points). After running the clustering 1000 times, a probability matrix for electrode pairs was defined as the ratio of numbers that electrode pairs clustered together, divided by the number of electrode pairs in the same random sampling for K-Means clustering. This ratio was calculated for all electrode pairs and used to create a matrix of pair-wise grouping probabilities. We then ran a second K-means clustering algorithm on this matrix to identify clusters that were similar in their network-level pair-wise grouping likelihood. Using a similar approach and random sampling of electrodes, we generated another pair-wise grouping probability matrix indicating how often electrodes were clustered together based on their pair-wise grouping likelihood (**Extended Data Fig. 4e**). Using this hierarchically defined pair-wise clustering likelihood, we ran a final K-means clustering on all electrodes, labeling them according to the probability of being stably grouped together for each cluster number. For each K (cluster number), we calculated clustering accuracy, confusion (probability of non-diagonal clusters), a confusion rank, and a ratio from dividing median accuracy (diagonals) by median confusion. We normalized each of the measures between 0 and 1 (1 showing the best, and 0 showing the worst performance between all clusters). We used these four measures and in a non-parametric vote, we chose the cluster number that outperformed others in this voting pool. This non-parametric measure shows how well each K performs to detect more stable clusters with high accuracy and low confusion level and estimate the optimal number of clusters. While there is no definitive answer on what number of clusters is the best, we selected the optimal number of clusters using this method to conduct further analysis. In summary, this clustering technique reduces clustering biases caused by outlier electrode pairs in all clustering realizations as well as stabilizes clusters.

To determine whether identified clusters were functionally specialized, we compared their responses evoked by cues and targets to their baseline activity levels. We calculated the averaged baseline normalized burst density within windows of 500 ms after cue onset and 750 ms after target onset. The Wilcoxon test was used to determine if this response was non-zero across the electrodes in each subnetwork. We corrected for multiple comparisons by using FDR for dependent samples with an alpha level of 0.05. Clusters that were significantly activated by cue events are referred to as “cue-subnetworks” and clusters that were significantly activated by target events are referred to as “target-subnetworks". We use “subnetworks” to indicate that these are functionally specialized modules within the larger brain-wide network engaged during attention. We consider the overarching network for each subject to be a network of all nodes. A subnetwork here is used to describe “modules” that we observe within the larger brain networks. The cue and target-subnetworks were found in each data set for the optimal number of clusters. For datasets, for which the optimal number of clusters did not contain distinct cue and target-subnetworks, if existed, we chose the next K (cluster number) with highest ranking in the clustering measures that contained both subnetworks (only in subject S12 we observed this).

### Time-lag analysis between cue and target-subnetworks

We calculated the HFAb-triggered-HFA for pairs of electrodes during the period between the target onset and the manual response to understand how the HFAbs were temporally ordered after the target onset in both the cue and target-subnetworks. Next, we compared HFAb-triggered HFA between electrodes in cue- and target-subnetworks to determine whether there was any asymmetry in the distribution of HFAbs between the two subnetworks. A total of six subjects (3 in experiment 1 and 3 in experiment 2) showed both stable cue and target-subnetworks. We then calculated once the HFAb-triggered HFA when HFA in the target-subnetwork was measured around the HFAbs in cue-subnetworks, and when the HFA in the cue-subnetwork was measured around the HFAbs in target-subnetworks. A lead state was indicated by HFAbs followed by stronger HFA power; a lag state was when HFAbs followed stronger HFA power.

First, we measured the maximum asymmetry around the HFAb onset across all subjects (by calculating the absolute difference between windows of varying lengths, 25 ms to 250 ms every 25 ms). We then calculated the averaged HFAb-triggered-HFA within 150 ms of HFAb onsets where asymmetry was at its maximum. Next, we asked if the value differed based on the directionality of the two subnetworks. For both directions, we used the Wilcoxon test to determine whether the asymmetry around HFAb onset is non-zero. Additionally, we tested whether HFAb-triggered HFA differs between the two directions after and before HFAbs.

To assess differences in HFAb-triggered HFA between cue- and target subnetworks, we used a non-parametric permutation testing approach. For each time bin (25ms) in the −500 to +500 ms window relative to HFAb onset, we calculated the observed difference in mean HFA between cue→target and target→cue electrode pairs. To test for significant differences between HFAb-triggered HFA in different directions, we used a permutation test with 10000 iterations, in which randomly shuffled electrode pair labels and generated a null distribution of mean differences. For each bin, a p-value was computed by comparing the observed difference in the two opposite directions, to the permutation distribution. To correct for multiple comparisons across time points, we used FDR correction for dependent samples at α = 0.05 (**Fig. 4a**). Additionally, cluster-based correction was implemented with a minimum cluster size threshold of 25 ms to identify temporally contiguous significant effects. Peak analysis was performed by identifying the maximum HFA amplitude and corresponding latency for each electrode pair after applying a 25 ms smoothing window (to avoid spurious peaks). Peak times are reported as mean ± standard error across all electrode pairs within each subject and task. We used Wilcoxon rank sum to test if the peak time distributions were significantly different in the opposite directions (**Fig. 4b**).

For each subject, we visualized the lead/lag interactions between stable cue and target-subnetworks. To achieve a better visualization of lead/lag interactions, we measured the HFA peak time for each electrode in cue/target-subnetworks relative to HFAb onset across all cluster numbers. Each electrode that was a member of the target-subnetwork or the cue-subnetwork was analyzed to determine its median peak-time-lag relative to the other cluster members. A median peak-time-lag of all electrodes satisfying this condition was then plotted, with red representing a lead, and blue representing a lag (**Fig. 4c**).

#### Delayed Mutual Information Analysis

We used mutual information (MI) which is a non-linear metric used in information theory to estimate the shared information between different time segments in two electrodes to find where their mutual predictability is maximized. For each electrode, we extracted data (HFAb density) from −1500 ms to 1500 ms around the target onset (and for the control analysis, around the cue onset). We normalized the data and calculated MI for segments of 750 ms sliding every 50 ms. The MI between electrodes X and Y is given by eq. 15:

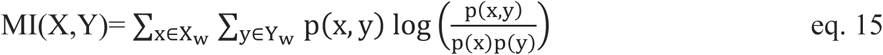

where p(x,y) is the joint probability distribution function of each windowed segment of electrodes X and Y, and p(x) and p(y) are their marginal probability distributions. We estimated the probability distribution for each variable and for the joint distributions using a histogram-based approach and binning the data (the number of bins was selected based on the Freedman-Diaconis rule ^88^ to balance the trade-off between estimation resolution and statistical reliability.

To further quantify how information is directionally coupled between cue and target-subnetworks, we calculated delayed mutual information (DMI). The DMI was calculated by comparing the temporal dynamics and dependencies between these two subnetworks (**Fig. 4d**).

The DMI between X and Y was calculated similar to MI, except that one of the timeseries was delayed incrementally to determine whether the past of one electrode is a better predictor of the future of the other electrode. The delay time-lag ranged from −500ms to 500ms with a step of 25ms, as in eq. 16.

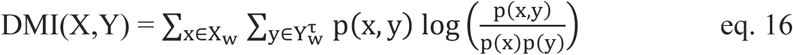

where X_w_ is segmented windows of electrode X, and 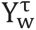 is segmented windows of electrode Y shifted by a time lag of τ.

The DMI analysis of each electrode pair can inform us about (i) when the electrodes showed maximum inter-predictability relative to an event onset (e.g., target), and (ii) at what time-lag the inter-predictability was maximized. To address this question, we extracted the DMI peaks in a 2D space, which gave us both the time-lag and the timepoint relative to the event onset where two electrodes showed maximum DMI values. We then asked if the time-lag of this peak is different between electrodes in the cue-subnetworks and electrodes in the target-subnetworks. For example, when we shifted the timeseries of electrodes in target-subnetworks, for each electrode in the cue-subnetwork, we measured its mean DMI relative to electrodes in target-subnetworks. We determined at what time lag and at what time relative to target onset DMI was maximized. We used the Wilcoxon test to see if the maximum DMI time-lag (τ_max_) occurred between electrodes in cue and target-subnetworks was non-zero. The lead and lag patterns in inter-predictability between cue and target-subnetworks were then determined by the sign of the average over τ_max_ for all electrodes in the shifting subnetworks.

### Modeling iEEG by spiking neural networks

We investigated high frequency activity in general and high frequency burst events in particular using a limited network of two recording sites placed on two interconnected networks of spiking neurons. For modeling the spiking networks, we used a framework described in^6^. Neuronal point-source models can accurately simulate electrical fields in cortical neural networks.^89^ Two cubic neuronal structures were simulated in three dimensions. Synaptic dynamics and connectivity patterns were implemented as explained in^6^. Each cubic network consisted of 1000 neurons (10 x 10 x 10, xyz). We modeled neurons as spherically symmetric points in a 3D grid (with r units on each axis). A recording disk was implemented on top of each network at a distance three times greater than the network depth (**Fig. 5a**). Pairwise connectivity was calculated based on anatomical studies (discussed below) and a distant-dependent Gaussian rule.^90,91^ The distant dependent connectivity factor is:

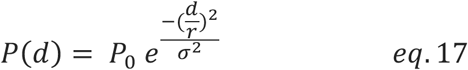

where r is the grid unit, *d* is the distance between two neurons, *σ* is the standard deviation of distances between neurons (Barral and Reyes, 2016), and *P*_0_ is a structural scaling factor (**Extended Data Fig. 8a**) reflecting the maximum connection probability across the network (see **Extended Data Table 2**).

Four interneuron types were used: Parvalbumin (PV), Calbindin (CB), Calretinin (CR), and Cholecystokinin (CCK) expressing interneurons. Excitatory neurons were divided into regular spiking neurons (70%), intrinsic low-threshold spiking bursting neurons (10%), and fast adapting regular spiking neurons (20%).

#### Neuron models

We used Izhikevich neuron model ^92^ for simulating regular spiking, burst spiking, fast spiking and low-threshold spiking neurons. Each neuron is modeled by a series of differential equations as in eq 18.

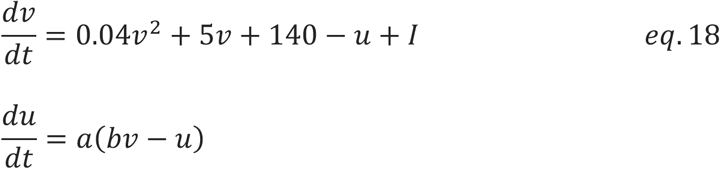

where variables *v* and *u* denote neural membrane potential and membrane recovery, respectively. Parameters *a* and *b* define the recovery rate and sub-threshold fluctuations sensitivity, respectively. For after-spike resenting the model uses two auxiliary equations as in eq. 19,

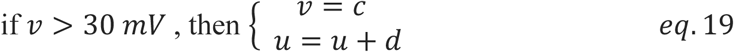

the parameter *c* resets the value of membrane potential *v* after a spike, and the parameter *d* adjusts the after-spike recovery variable *u*. Parameters for different neuron types were chosen as suggested in ^93^ to approximately generate firing patterns of each neuron type (See **Extended Data Table 2** for each neuron type parameter).

#### Neural connectivity

Besides distance-based connectivity factors, neuron types had different connection probabilities. We adapted the scaling factor for connectivity between excitatory neurons and inhibitory interneurons from ^91^. Additionally, rodent anatomical studies were considered in determining the connectivity among different types of neurons. Generally, PV interneurons inhibit themselves and VIP interneurons (likely similar to CR), whereas SOM interneurons (likely similar to CB) do not inhibit each other, and VIP interneurons disinhibit SOM interneurons preferentially ^94,95^. The connectivity between networks was defined by a probability and a rate of connection. This connectivity was attributed primarily to excitatory neurons (90%).

#### Simulation of post-synaptic potentials

For each neuron the postsynaptic potentials were modeled by a biexponential function as in eq. 20,

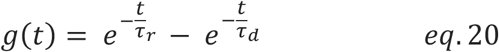

where *τ*_*r*_ and *τ*_*d*_ denote the rise and decay time constant of postsynaptic current, respectively. The biexponential function was implemented through a second-order ordinary differential equation as in eq. 21.

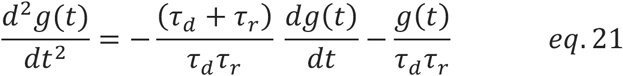

Where t is time relative to spike, and g(t) is the synaptic conductivity. The rise time was set similarly to ∼1ms for all neurons while the decay time for pyramidal neurons were and interneurons varied from ∼6ms to 24ms^96–99^ as shown in **Extended Data Table 2**.

#### Network external inputs

Each network was fed external currents to generate firing rates similar to cortical neurons. Each neuron received a cosine input. The objective was to first control externally induced rhythmicity in network activation by frequency, as well as phase coherence between the input function and neurons. On average, each network received external input sufficient to generate a 5 Hz firing rate.^100^ The external input to pyramidal neurons was three times greater than the external input to inhibitory neurons. In addition, a Brownian noise 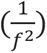 was added to the input as per previous experimental observations.^101^

For nonrhythmic input, we considered slow and ultraslow oscillatory (<1 Hz) input to each neuron with an initial phase lag. Then a coherence index was used to determine the distribution of phase lags among neurons. Thus, the phase range was defined:

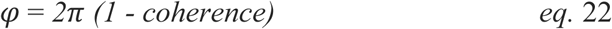

For nonrhythmic input, we ran 100 simulations in which coherence varied from 0 to 1 with a step size of 0.01 (**Fig. 5e**). For Networks with reciprocal connections, we fed the networks with uncorrelated inputs, but both networks were reciprocally connected (**Fig. 5g**). The connectivity strength was changed by one order of magnitude from 0.001 to 0.1 and the simulation was run 100 times for each value. For networks with feedforward connection, we simulated feedforward networks in which the connections between the two networks are unidirectional. The connectivity strength was changed from 0.001 to 0.1 by one order of magnitude (**Fig. 5h**). For the rhythmic input, we incrementally increased the cosine input frequency up to 12 Hz for each simulation run (**Fig. 5i**). For networks with shared input, we ran 100 simulations in which networks were not connected but received correlated inputs (5-25 percent of neurons received the same input, not shown).

To investigate the effect of an external stimuli on directional networks, we simulate a feedforward network as explained before with an internetwork connectivity ratio of 9 to 1 (network 1 and 2 respectively, **Extended Data Fig. 8e**). We then fed network 1 with an external input pulse of 500 ms duration and 250 ms duty cycles (**Extended Data Fig. 8f**). The input was fed to 50 percent of neurons in the network 1. We then calculated the time resolved PLV in both networks as explained before (**Fig. 5f**, **Extended Data Fig. 8g**).

#### Integration of postsynaptic currents and neural activities at the recording sites

For each simulation, we estimated the field potential at two recording sites (black disks **Fig. 5a**). By assuming neurons as point-source field, we measured field dynamics at each recording site by summing the attenuated membrane potentials and the postsynaptic currents from all neurons. We estimated the voltages at the recording disk by considering membrane and post-synaptic potentials as electrical dipoles (with negligible distance between poles relative to the recording disk). The electric filed is then represented by *E*(*r*, *t*), where *r* and *t* denote the distance from the source and the time, respectively. The extracellular potential is then calculated by:

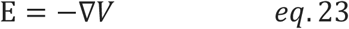

Using Ohm’s law, the electric field at distance r from each dipole with a current density of *I*_*n*_ (t) is equal to:

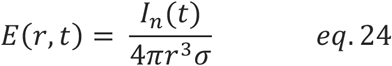

with *I*_*n*_(*t*) represented as:

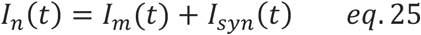

where *σ* denotes the medium conductivity (which we assumed is independent of distance from sources). *I*_*m*_(*t*) and *I*_*syn*_(*t*) are the transmembrane current and the synaptic current of each neuron respectively. By integrating the electric field, we calculated the potential at the disk by:

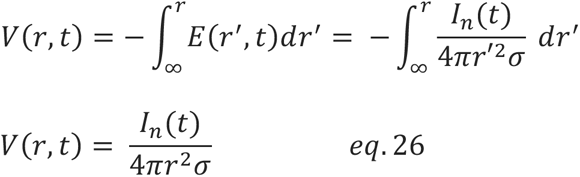

The voltage recorded at each site is calculated as in eq 27:

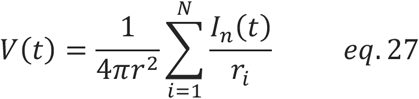

where *r*_*i*_ is the distance between the recording disk and the *i^th^* point-source, and N is equal to the total number of neurons in each network.

We used a non-ohmic filter to attenuate higher frequencies by getting insights from.^102^ We implemented an exponential attenuation in the frequency domain. First for each signal we computed the FFT. We then applied an exponential attenuation factor to the magnitude of the frequency components (**Extended Data Fig. 8b**). This factor exponentially decreases the amplitude for higher frequencies as in eq. 28:

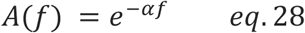

Where *A*(*f*) is the attenuation factor for frequency *f, and α* is a parameter that controls the rate of exponential decay which we set to 0.01. After multiplying the attenuation factor, we performed the inverse FFT (iFFT) to transform the signal back to the time domain.

For each condition, we ran simulations 100 times, each simulating iEEG signals and neural activity for three seconds. We detected HFAbs at each recording site and calculated the amplitude of the analytical signal as explained in (eq. 1). Next, we examined whether there was a correlation between HFAb density and aggregated spike density in each network. The spike density was calculated and smoothed using a Gaussian window of 25 ms. We measured the correlation coefficient between aggregated spike density and burst density for each simulation. On average, HFAb events were significantly correlated with spike densities in each network.

In a control condition, we randomly assigned burst times and examined the correlation between burst density and spike density. This randomization was performed 1000 times and we found the 95% confidence interval under the null hypothesis that burst density is not related to spike density. All other analyses of the modeling results were conducted in a similar manner to those of the experimental data.

#### Four-Network Model of Spatial Attention

We extended our model to a four-network architecture mimicking the cue-target processing hierarchy (**Fig. 5j**). The model contained two spatially-selective cue networks (N1, N2) and two target networks (N3, N4), with dominant feedforward connections from cue to target networks (5 times stronger than feedback).^103^ Intra-network connection strength was set 2 times greater than the inter-network connectivity strength.^104^ To simulate the spatial attention task, we implemented three distinct external input functions:

##### Trial initialization

A decaying exponential input (τ = 4s) was delivered to all networks at trial onset, decreasing to 25% of initial amplitude by trial end simulating baseline states maintained throughout the trial.

##### Cue processing

Upon cue presentation, one cue network received spatially-selective input modeled as a double-exponential function (*τ_rise_* = 50ms, *τ_decay_* = 1s).^56,105^

##### Target processing

Target networks received double-exponential input (*τ_rise_* = 100ms, τ = 500ms) at target onset. In valid trials, where targets appeared at cued locations, the previously activated cue network received additional weak reactivation at 50% of original cue amplitude, simulating either feedback or sustained attention signals from frontal-parietal networks or inputs from subcortical structures such as pulvinar.^56,105,106^

We simulated 150 trials for each condition (valid and invalid cues) with 4s trial duration. HFAbs were detected using identical parameters as empirical data, and we calculated HFAb responses in each network and delayed mutual information (DMI) between cue-target network pairs.

#### HFAb-based detection modelling

To link burst dynamics to behavioral performance, we extracted HFAb rates from target networks within 500ms windows following target onset for both valid and invalid conditions. Detection probability was modeled using a logistic function:

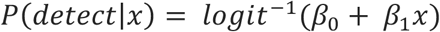

where *x* represents HFAb rate, and *β₀, β₁* are model parameters estimated via maximum likelihood. We fitted detection thresholds to invalid trials to achieve ∼85% accuracy (matching empirical performance), then evaluated whether valid trials exceeded this threshold. In addition, HFAb rates were binned into 100 bins spanning the full range to calculate detection probabilities for each condition (**Fig. 5m**).

## Data Availability

Preprocessed data used for generating these results are deposited at Figshare and will be accessible through https://doi.org/10.6084/m9.figshare.30434683.107 Source data are provided with this paper.

## Code Availability

Custom programming codes for analysis and modeling are available from https://github.com/banaiek/Attentional_Routing/.

