## Supplementary Information for "High frequency bursts facilitate fast communication for human spatial attention"

### Supplementary Materials

**Extended Data Figs.** 1 to 8
**Supplementary Tables** 1 to 2

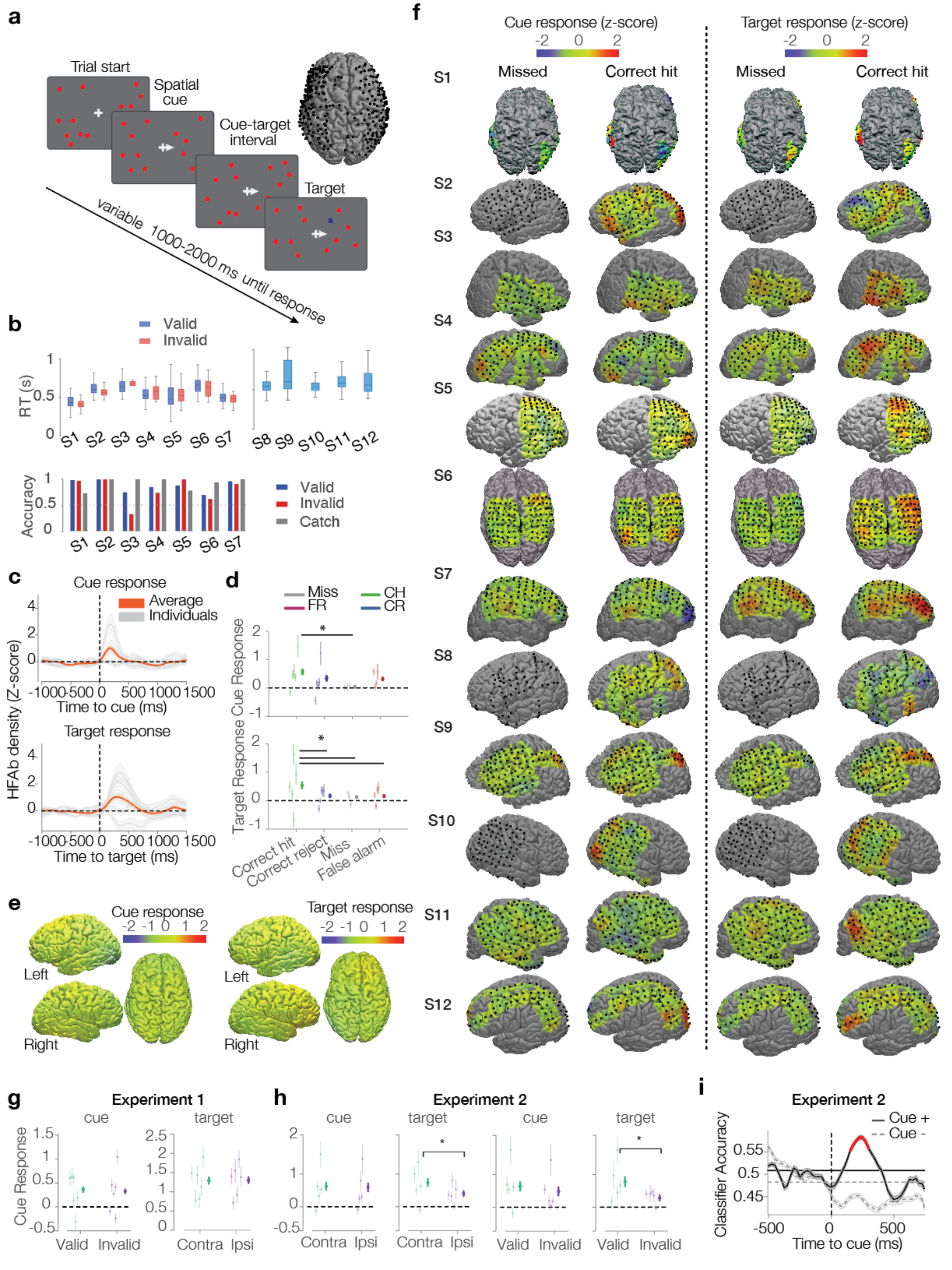
a
Trial start
Spatial cue
Cue-target interval
Target
variable 1000-2000 ms until response
b
Valid
Invalid
RT (s)
S1 S2 S3 S4 S5 S6 S7 S8 S9 S10 S11 S12
Accuracy
Valid
Invalid
Catch
c
Cue response
Average
Individuals
HFAb density (Z-score)
Time to cue (ms)
Target response
Time to target (ms)
d
Miss
FR
CH
CR
Cue Response
Target Response
Correct hit
Correct reject
Miss
False alarm
e
Cue response
Target response
Left
Right
f
Cue response (z-score)
Target response (z-score)
Missed
Correct hit
S1 S2 S3 S4 S5 S6 S7 S8 S9 S10 S11 S12
g
Experiment 1
cue
target
Cue Response
Valid
Invalid
Contra
Ipsi
h
Experiment 2
cue
target
Contra
Ipsi
Valid
Invalid
i
Experiment 2
Cue +
Cue -
Classifier Accuracy
Time to cue (ms)

**Extended Data Fig. 1. Behavioral performance and HFAb activation patterns across experiments.** **a**, Task structure of experiment 2. Subjects hold their gaze fixation to the center of a screen (a white plus sign) with red circles turning on and off. A spatial cue endogenously cues subject's attention to a hemifield. A target appears at one hemifield and subjects should report whether the target was seen in the cued hemifield. The brain shows the localization of electrodes across all subjects. **b**, Reaction times (top) and accuracy (bottom) for individual subjects across valid, invalid, and catch trials (trials with no targets in experiment 1). **c**, Population-averaged HFAb density around cue (top) and target (bottom) onsets, similar to **Fig. 1b** in experiment 2. **d**, HFAb responses grouped by trial outcomes (correct hit, correct reject, miss, false alarm) for cue (top) and target (bottom) epochs. Asterisks indicate significant differences (Kruskal-Wallis test $p = 0.003$ for cue, and $p < 0.001$ for target response; followed by Dunn's test, *$p < 0.05$). **e**, Group topography of cue (left) and target (right) response for incorrect trials. **f**, Individual subject topographies for correct and incorrect trials, similar to **Fig. 1h**. **g,h**, HFAb responses comparing valid/invalid cues and contra/ipsi laterality conditions for experiments 1 (**g**) and 2 (**h**), similar to **Fig. 1i,j**. Asterisks indicate significant main effect (GLME, $p<0.05$). **i**, Classifier accuracy predicting trial outcomes for cue-responsive (Cue+) and cue-unresponsive (Cue-) in experiment 2. Red line indicates significant prediction above baseline and chance (binomial test, $p < 0.05$, FDR corrected).

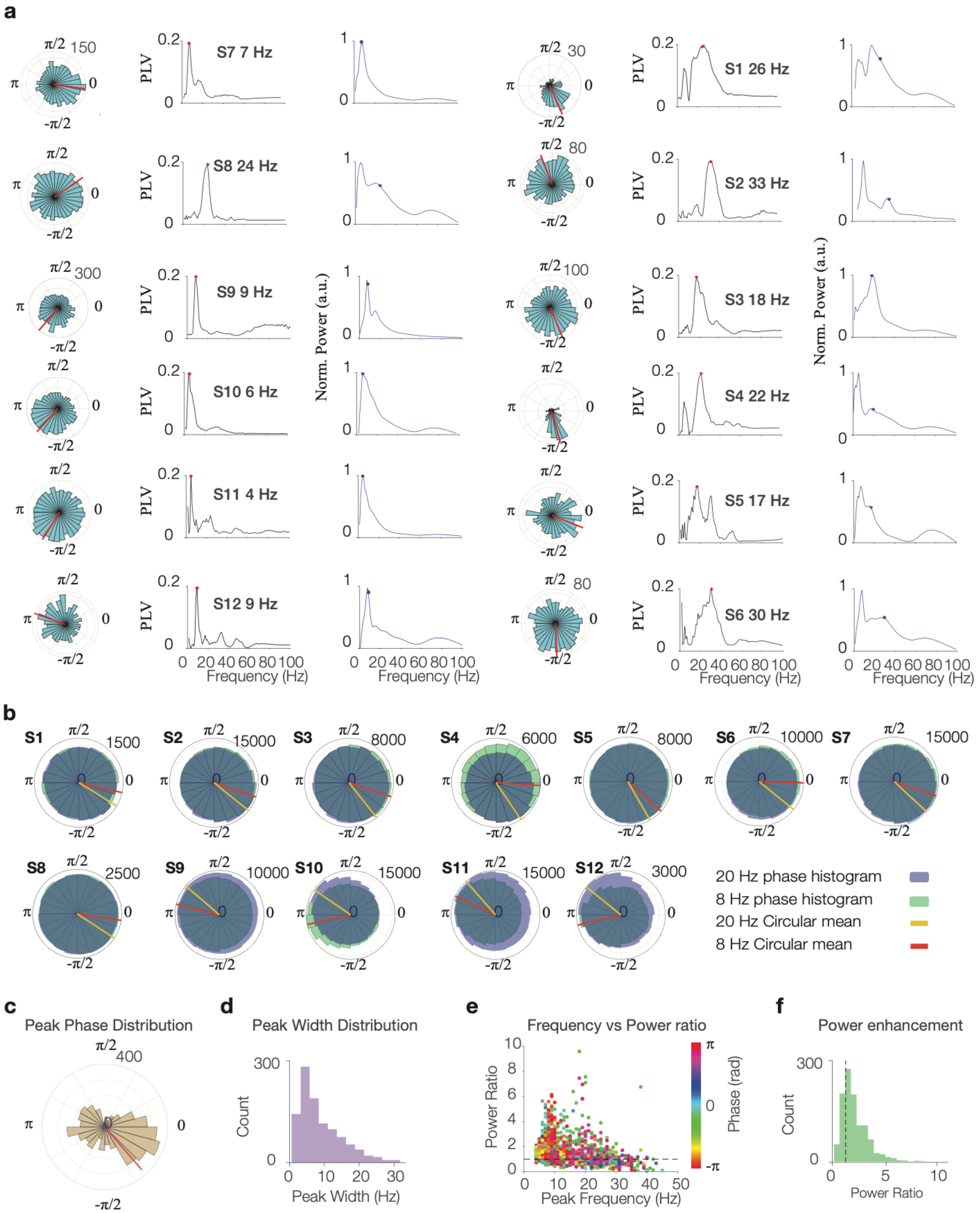


**Extended Data Fig. 2. HFAb phase-locking to low-frequency rhythms across subjects. a**, Individual examples showing HFAb phase distributions at the peak (left), phase-locking value (PLV, middle), and HFAb-triggered spectrum (right) from different subjects. Peak frequencies noted for each example in red and blue circles. **b**, Phase histograms of HFAbs at theta/alpha (8 Hz, purple) and beta (20 Hz, green) frequencies for all subjects. Orange and red lines indicate circular means. **c**, Distribution of peak phases across all electrodes. **d**, Distribution of phase-locking peak widths (mean: 7.3 ± 0.3 Hz). **e**, Relationship between peak frequency and power ratio (HFAb-triggered/baseline), colored by phase value. **f**, Distribution of power enhancement ratios at phase-locking frequencies (median: 2.1-fold increase), dashed line shows 1.

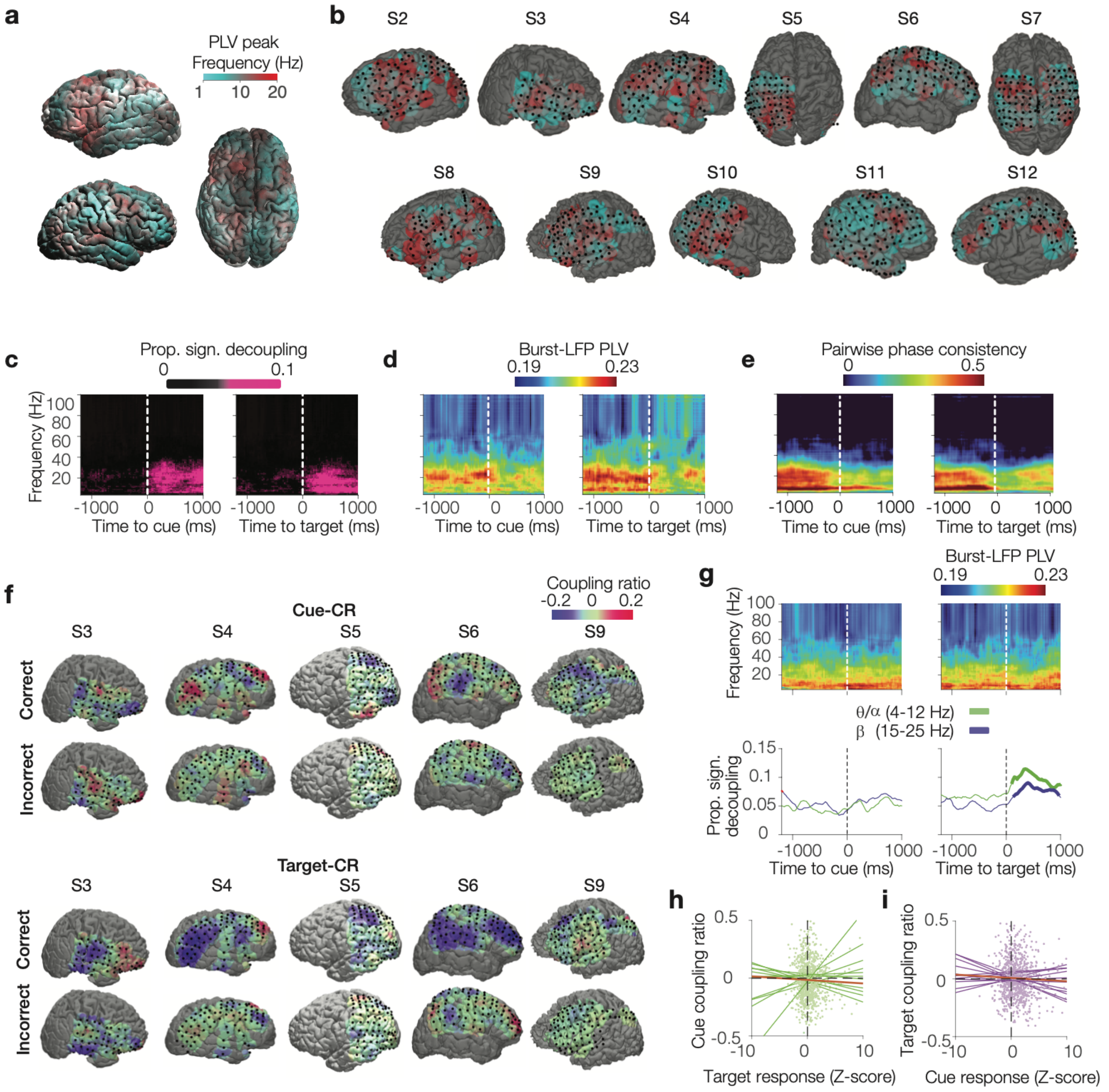


**Extended Data Fig. 3. HFAb synchronization with low frequency activity in experiment 1 and 2. a,** Group-average spatial pattern of observed frequency peak of HFAb phase locking to LFP. **b**, Individual examples showing the spatial pattern of the observed frequency peaks of the HFAb phase locking to LFP. **c**, Corresponding to **Fig. 2e**, showing the proportion of time-frequency points where phase locking was significantly lower than baseline ($P < 0.05$, random permutation test). **d**, Similar to **Fig. 2e**, but after removing event-related potential from the LFP (see **Methods**). **e**, Group-average heatmaps of pairwise phase consistency (PPC) as a measure for HFAb synchronization to LFPs, Similar to **Fig. 2e**. **f**, Examples of the coupling ratio between HFAb and low frequency (4-25 Hz) LFP following cue and target onsets in correct and incorrect trials. **g**, Similar to **Fig. 2e** for experiment 2. **h,i**, Regression plots showing correlation of coupling ratios following cue onset with target responses (green, **h**), and coupling ratios following target onset with cue responses (purple, i). Scatter points denote electrodes, lines indicate individual subjects with orange line showing the regression across all subjects.

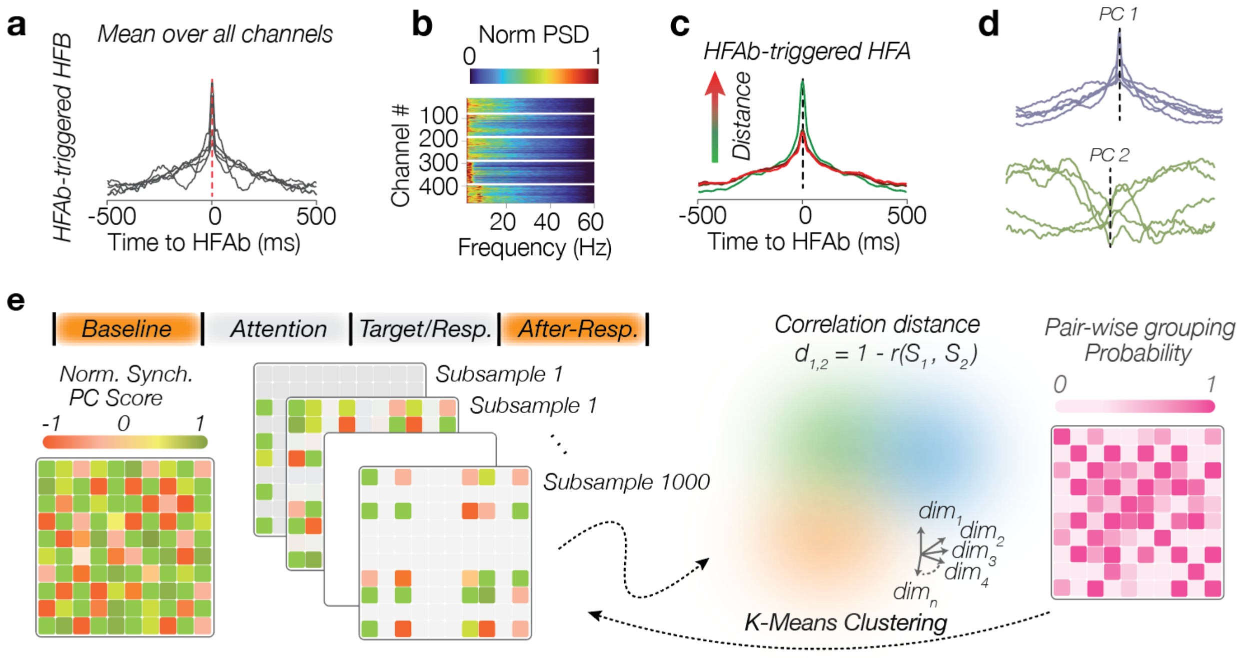


**Extended Data Fig. 4. A network clustering approach based on the synchronization of HFAbs between electrodes. a,** The average HFAb-triggered HFA for individuals in experiment 2. **b**, The normalized PSD for HFAb-triggered HFA in experiment 2**. c,** HFAb-triggered HFA for electrodes distanced in 4 different quantiles (25,50, 75, 100 mm), ranging from green (short) to red (long) in experiment 2, similar to **Fig. 3c**. **d**, The first and second principal components of HFAb-triggered HFA for individual subjects in experiment 2. **e**, A schematic demonstration of network clustering algorithm. We used HFAbs outside of cue/delay and target/response periods. The network synchrony matrix shows the loading values for each electrode pair on the synchronized component. A K-means clustering was performed on randomly selected electrode samples for different cluster numbers (K = 2 to 8). We calculated a pair-wise grouping probability matrix in which each element indicates how likely it is that two electrodes will be grouped together. The next step was clustering with network subsampling, similar to the previous step but based on the pairwise grouping likelihood matrix. The final clustering of the pair-wise grouping likelihood results indicated stable clusters for each K (see **Methods**).

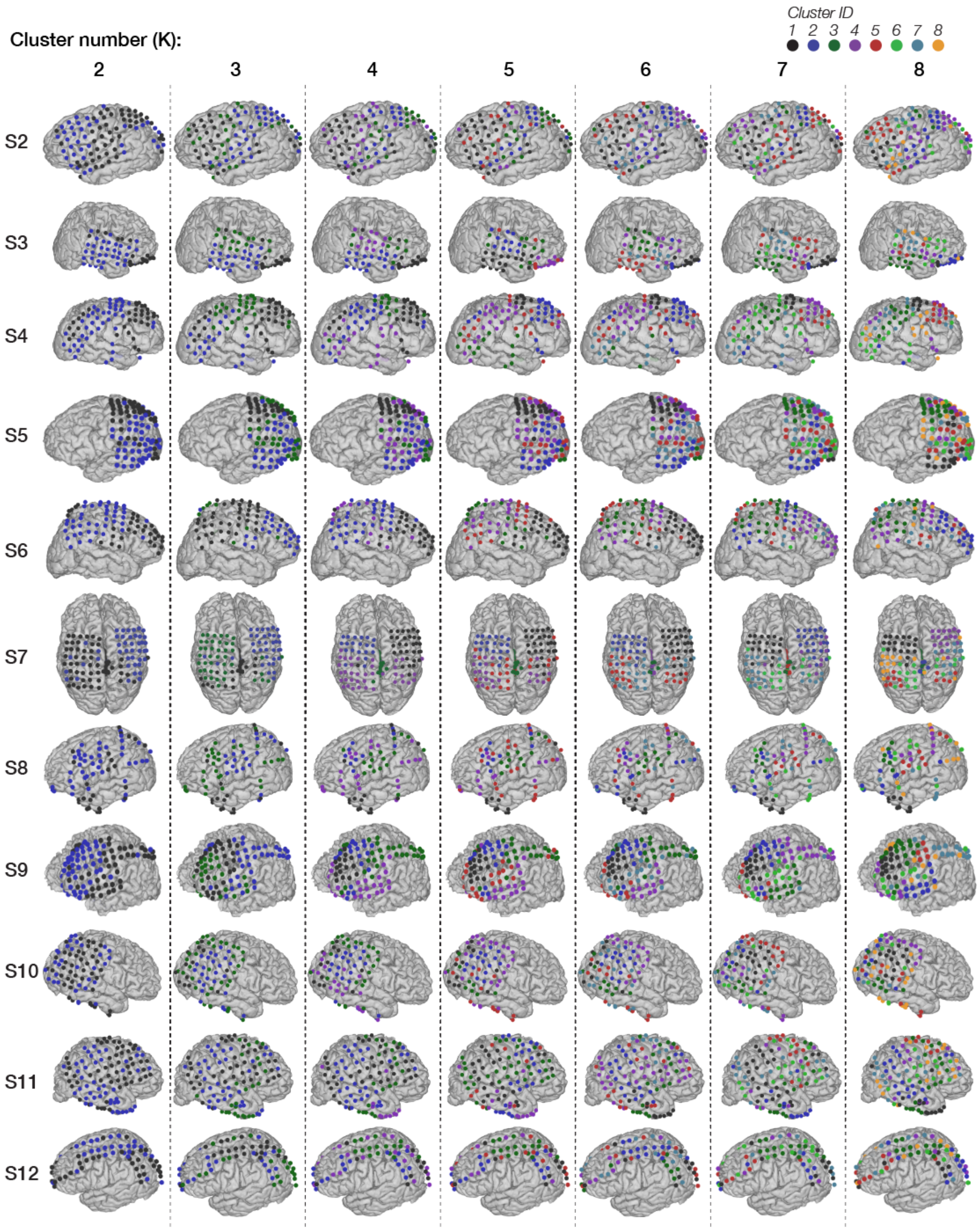


**Extended Data Fig. 5. Organization of clusters based on cluster numbers.** Columns from left to right show the results for cluster numbers K = 2 - 8. The cluster IDs are sorted by cluster stability.

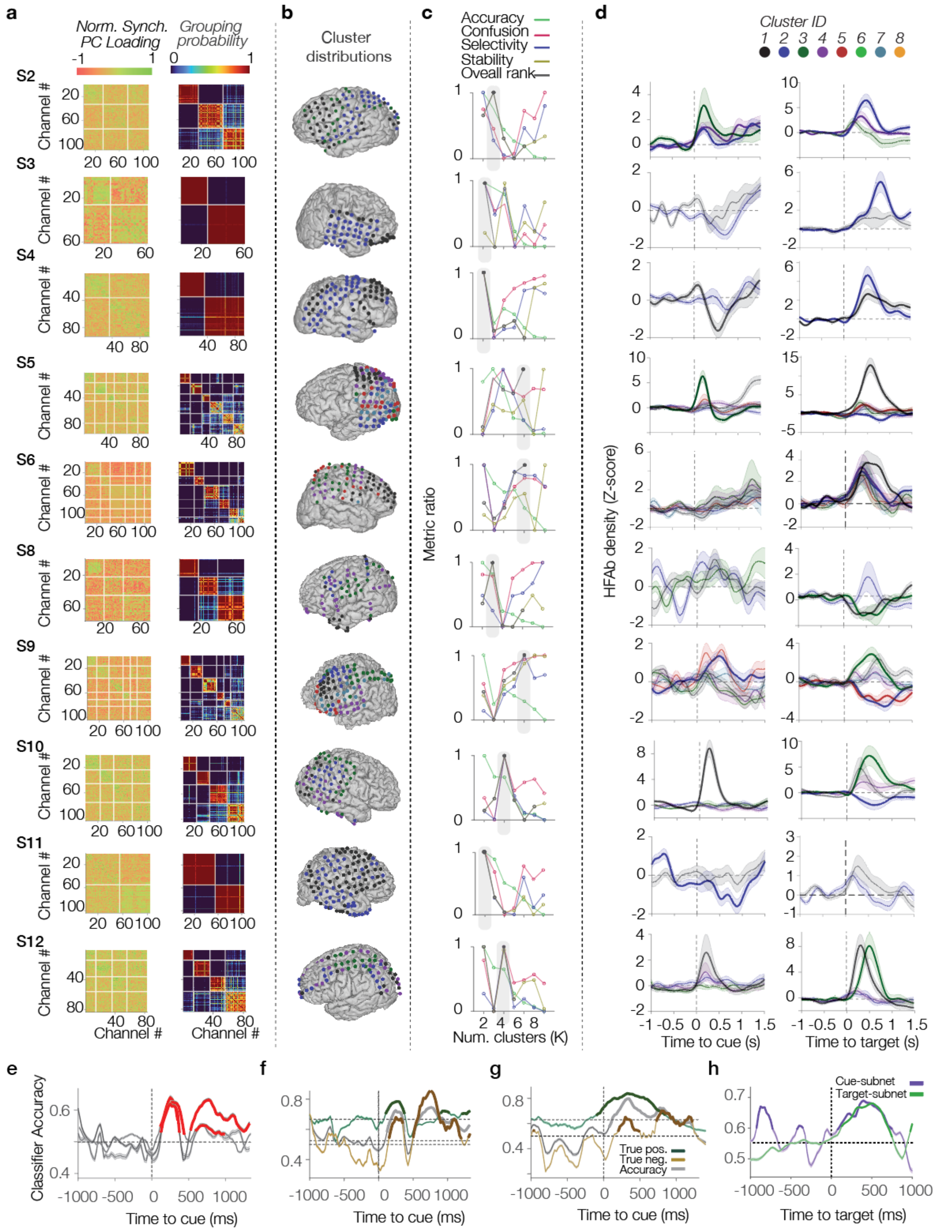


**Extended Data Fig. 6. Clustering results for individual subjects. a**, Color-coded loading values on the synchronized PC (left) and the pairwise grouping probability (right) for each subject. **b**,

Electrode distributions for optimal cluster numbers. **c**, Cluster number selection using four metrics with nonparametric voting rank (black). The accuracy is determined by the median diagonals, the confusion by the median nondiagonal, selectivity by the relative rank of the diagonal over the highest nondiagonal rank, and stability by the relative rank of the diagonal over the nondiagonal rank. **d**, HFAb density around cue and target onsets for each cluster, similar to **Fig. 3f**, across subjects. Shaded error bars indicate the standard error of the means, thicker lines indicate significant functional subnetworks. **e,** Leave-one-subject-out classifier analysis, similar to **Fig. 3h**. **f, g**, Classifier accuracy predicting correct trials (green), errors (brown), and overall accuracy (gray) for (**f**) experiment 1 and (**g**) experiment 2. **h,** outcome prediction by cue- and target-subnetworks following target onset. Shaded error bars show standard error of the mean. Thick lines indicate timepoints above baseline and chance ($P < 0.05$, binomial test, FDR corrected).

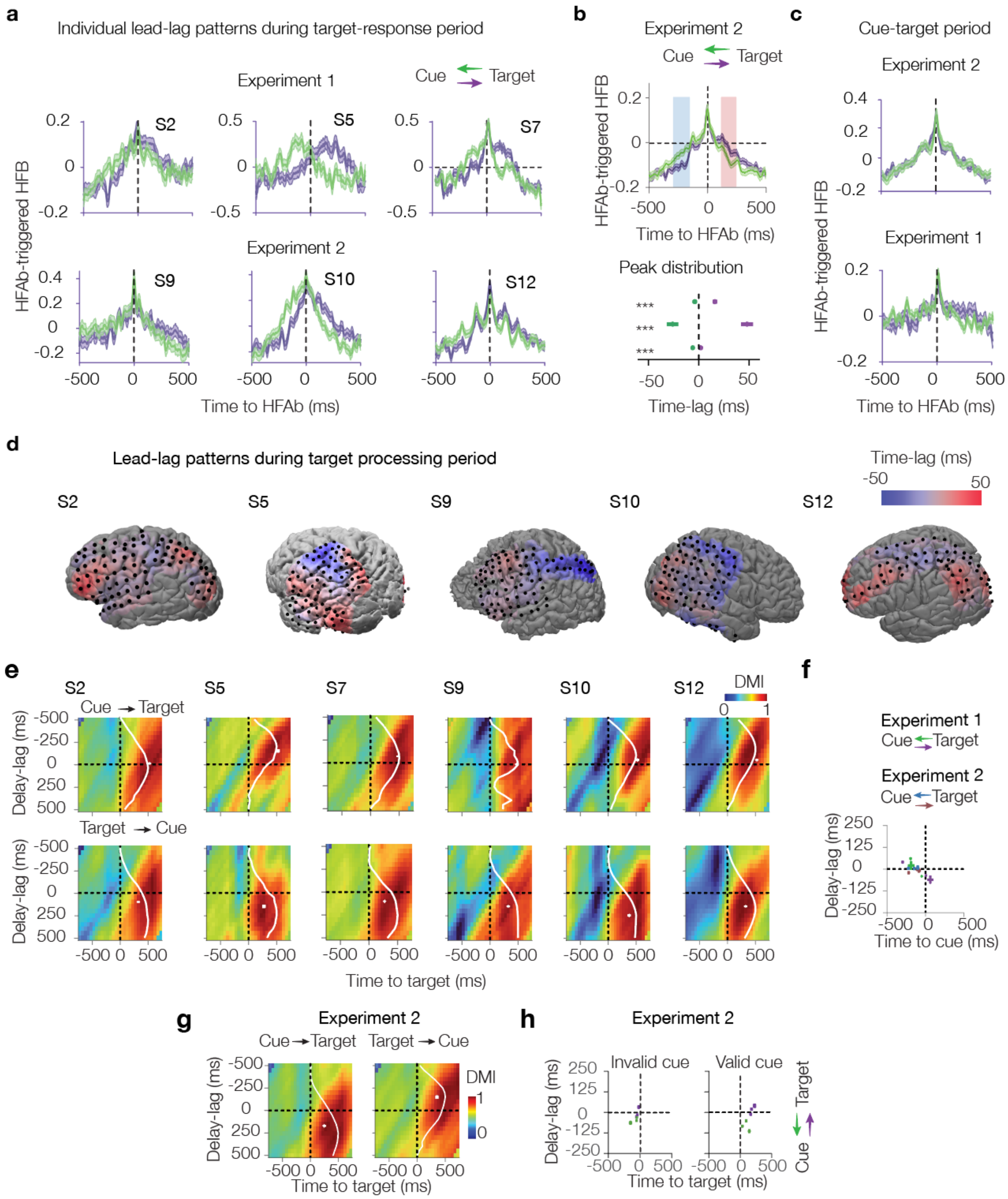


**Extended Data Fig. 7. Temporal precession between cue and target-subnetworks. a**, HFAb-triggered HFA examples for individual subjects, similar to **Fig. 4a**. **b**, Top: Group-level average of HFAb-triggered HFA in experiment 2, similar to **Fig. 4b**. Shaded regions indicate significant lead-lag patterns (permutation test, $p < 0.05$; red: cue leads, [135-244] ms, blue: target leads, [-298 to -155] ms). Bottom: Peak time-lag distributions in experiment 2 (mean: 14.9 ± 4.7 ms) **c**, HFAb-triggered HFA during cue/delay period, Similar to **Fig. 4a. d,** Lead-lag pattern topography for individual subjects, similar to **Fig. 4c. e**, DMI heatmaps for individual subjects, similar to **Fig. 4d**. **f**, DMI analysis around cue onset, similar to **Fig. 4e**. **g,** Group-average DMI in experiment 2, similar to **Fig. 4d.** DMI peaked 297.7 ± 19.6 ms after target onset with 125.6 ± 26.2 ms time-lag ($n = 3$). **h**, DMI peak distributions for invalid/valid trials in experiment 2, similar to **Fig. 4e.**

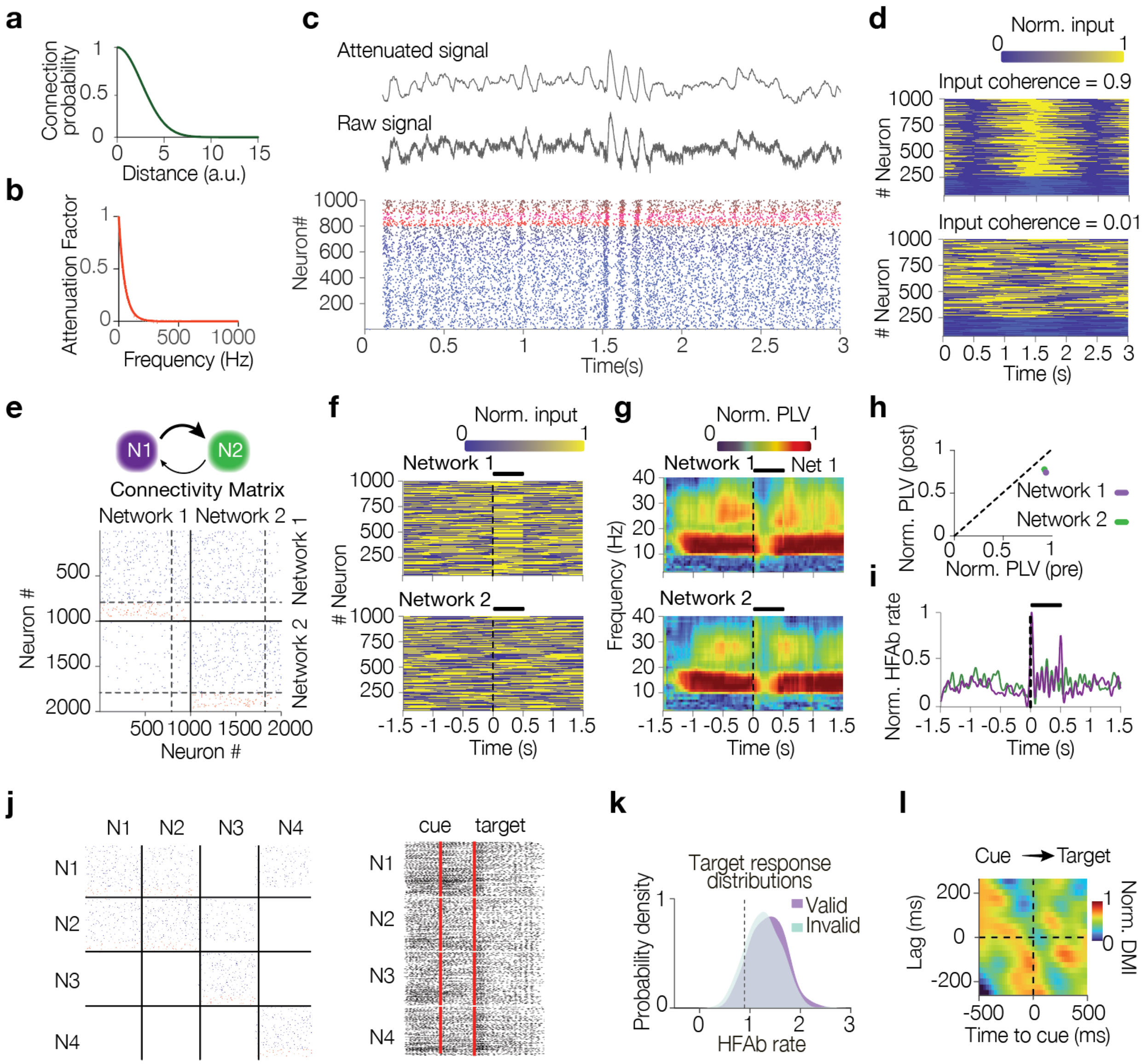


**Extended Data Fig. 8. Computational modeling of iEEG signal. a, Connection probability between neurons decreases with distance. b, Frequency-dependent attenuation factor for iEEG signals at the recording disks. c**, A complete trial simulation example. Raster shows the activity of neurons in one network (bottom, blue and red show excitatory and inhibitory neurons, respectively). Raw and attenuated traces correspond to field dynamics of the same network. **d, Network activity with coherent (top) and random (bottom) input phases. e**, **Connectivity matrix for feedforward** network **(N1→N2,** each scatter point indicates whether two neurons have excitatory (blue) or inhibitory (red) connections**). f**, An input design evaluating how a transient stimulus affects HFAb coherence with LFP in a network as shown in E (network 1 (top) receives an impulse input). **g**, PLV changes in both networks following transient input, similar to **Fig. 5g**. **h** In both networks 1 and 2, the PLV drops within 500ms of stimulus onset (P < 0.001, Wilcoxon test). **I,** Normalized HFAb rate relative to stimulus onset. **j,** Four-network architecture and example raster activity during spatial attention task simulation. **k**, HFAb rate distributions in target networks for valid versus invalid trials. **l**, Normalized DMI between cue and target networks following cue onset, similar to **Fig. 5l**.

### Supplementary Tables

| Subject ID | Cue + | Target + | Cue Subnetworks | Target Subnetworks |
|---|---|---|---|---|
| **S1** | IFG, TPJ, V3b, hV4 | ISP0, MFG, MTG, TPJ | | |
| **S2** | Cingulate Gyrus, ISP2, IFG, IPL, LO2, MFG, Postcentral Gyrus, SFG, | Cingulate Gyrus, ISP1, ISP3, IPL, MFG, Paracentral Lobule, Postcentral Gyrus, Precentral Gyrus, SFG, STG | Cingulate Gyrus, ISP2, ISP3, IFG, IPL, MTG, Paracentral Lobule, Postcentral Gyrus, Precentral Gyrus, STG | Cingulate Gyrus, ISP1, ISP2, ISP3, IFG, IPL, MFG, MTG, Paracentral Lobule, Postcentral Gyrus, Precentral Gyrus, SFG, STG |
| **S3** | IPL | IFG, IPL, ITG, MTG, Orbital Gyrus, Precentral Gyrus, SFG, STG, TPJ | | IFG, IPL, ITG, MFG, MTG, Postcentral Gyrus, Precentral Gyrus, STG, TPJ |
| **S4** | IPL, MFG, hMT, Postcentral Gyrus, LO2 | Angular Gyrus, FEF, FFG, IFG, IPL, MFG, Postcentral Gyrus, Precentral Gyrus, SFG, STG, TPJ, | | Angular Gyrus, Anterior Cingulate, Cingulate Gyrus, FEF, FFG, IFG, IOG, IPL, ITG, MFG, MTG, PHG, Postcentral Gyrus, Precentral Gyrus, SFG, STG, TPJ |
| **S5** | FFG, ISP2, ISP3, LO2, MOG, MTG, Postcentral Gyrus, hMT | FEF, FFG, IPL, MTG, Postcentral Gyrus, Precentral Gyrus, SFG, TPJ | FFG, LO2, ISP3, MOG, Postcentral Gyrus, hMT, Postcentral Gyrus | FEF, IPL, Postcentral Gyrus, Precentral Gyrus, SFG |
| **S6** | FEF, IPL, Postcentral Gyrus, SFG, TPJ | FEF, IPL, MFG, Paracentral Lobule, Postcentral Gyrus, Precentral Gyrus, Precuneus, SFG, STG, TPJ | | FEF, ISP2, ISP3, IPL, MFG, Paracentral Lobule, Precentral Gyrus, Precuneus, SFG, STG, TPJ |
| **S7** | ISP2, ISP3, IPL, Postcentral Gyrus, SFG, SPL, TPJ | FEF, ISP2, ISP3, ISP5, IPL, Paracentral Lobule, Postcentral Gyrus, Precentral Gyrus, Precuneus, SFG, SPL, TPJ | ISP2, ISP3, ISP5, IPL, SPL, TPJ, Postcentral Gyrus, | FEF, ISP2, ISP3, ISP5, IPL, Paracentral Lobule, Postcentral Gyrus, Precentral Gyrus, Precuneus, SPL, SFG, TPJ |
| **S8** | LO1 | FEF, ISP1, ISP2, MFG | | FEF, ISP1, ISP2, MFG |
| **S9** | ISP2, ISP3, MFG, STG | ISP1, ISP2, ISP3, MFG, MTG, Precentral Gyrus, SFG, STG | ISP1, ISP2, ISP3, MFG, STG | ISP1, ISP2, ISP3, MFG, MTG, Precentral Gyrus, SFG, STG |
| **S10** | ISP2, ISP3, MFG, Postcentral Gyrus, FEF | FEF, ISP2, ISP3, IFG, IPL, MFG, MTG, Postcentral Gyrus, Precentral Gyrus, SFG, STG | FEF, ISP2, IPL, MFG, Postcentral Gyrus, Precentral Gyrus, SFG, STG | FEF, ISP2, IFG, IPL, MFG, MTG, Postcentral Gyrus, Precentral Gyrus, SFG, STG, TPJ, V3b |
| **S11** | ISP2, MFG, hV4 | ISP1, ISP2, ISP3, IFG, MFG, MTG, Postcentral Gyrus, STG | | |
| **S12** | ISP2, ISP3, Postcentral Gyrus | FEF, ISP1, ISP2, ISP3, IPL, MFG, MTG, Postcentral Gyrus, Precentral Gyrus, SFG | ISP1, ISP2, ISP3, Postcentral Gyrus, MFG, Precentral Gyrus, | FEF, ISP1, ISP2, ISP3, IPL, MFG, MTG, Postcentral Gyrus, Precentral Gyrus, SFG |

**Supplementary Table 1.** List of brain areas containing electrodes that showed significant HFAb response to cue (cue +) and target (target +, see **Methods**), as well as electrodes in cue- and target-activated subnetworks. The abbreviations are: Inferior Frontal Gyrus (IFG), Temporoparietal Junction (TPJ), Visual area 3b (V3b), human Visual area 4 (hV4), Intraparietal Sulcus (IPS1, IPS2, IPS3, IPS5), Middle Frontal Gyrus (MFG), Middle Temporal Gyrus (MTG), Inferior Parietal Lobule (IPL), Lateral Occipital area 2 (LO2), Superior Frontal Gyrus (SFG), Superior Temporal Gyrus (STG), Inferior Temporal Gyrus (ITG), Frontal Eye Field (FEF), Fusiform Gyrus (FFG), Inferior Occipital Gyrus (IOG), Parahippocampal Gyrus (PHG), Middle Occipital Gyrus (MOG), human Middle Temporal/V5 (hMT), Superior Parietal Lobule (SPL), and Lateral Occipital area 1 (LO1).

| Cell-type<br><br>Parameters | Pyr | PV | CCK | CB | CR |
|---|---|---|---|---|---|
| **population** | 0.76 | 0.07 | 0.02 | 0.9 | 0.06 |
| **a** | 0.02 | 0.1 | 0.05 | 0.02 | 0.02 |
| **b** | 0.2 | 0.23 | 0.23 | 0.23 | 0.23 |
| **c** | -65 | -65 | -65 | -65 | -65 |
| **d** | 8 | 2 | 2 | 2 | 2 |
| $\boldsymbol{\tau_r}$ | 1 | 1 | 1 | 1 | 1 |
| $\boldsymbol{\tau_d}$ | 6.4 | 8 | 12.4 | 16 | 16 |
| **Pyr** | 0.3 | 0.5 | 0.5 | 0.5 | 0.5 |
| **PV** | 0.6 | 0.4 | 0.25 | 0.15 | 0.05 |
| **CCK** | 0.6 | 0.25 | 0.4 | 0.15 | 0.05 |
| **CB** | 0.6 | 0.6 | 0.6 | 0.05 | 0.25 |
| **CR** | 0.05 | 0.05 | 0.05 | 0.6 | 0.05 |

**Supplementary Table 2.** Parameters used for modeling different neuron types. The parameter $a$ indicates a recovery rate variable, $b$ the sensitivity to sub-threshold fluctuations, $c$ the membrane potential, $d$ adjusts the after-spike recovery variable, $\tau_r$ synaptic potential rise time, and $\tau_d$ synaptic potential decay time (see **Methods**). The last 5 rows indicate the connectivity weight matrix between different cell types.
